# Impaired spatial coding and neuronal hyperactivity in the medial entorhinal cortex of aged *App^NL-G-F^* mice

**DOI:** 10.1101/2024.11.26.624990

**Authors:** Gustavo A. Rodriguez, Eva F. Rothenberg, C. Oliver Shetler, Andrew Aoun, Lorenzo Posani, Srujan V. Vajram, Thomas Tedesco, Stefano Fusi, S. Abid Hussaini

**Author notes:** Corresponding author: S. Abid Hussaini, PhD, Assistant Professor, Department of Pathology & Cell Biology, Taub Institute for Research on Alzheimer’s disease and the Aging Brain Columbia University Irving Medical Center, P&S 12-420C, 630 W 168th St New York, NY 10032.

## Abstract

The progressive accumulation of amyloid beta (Aβ) pathology in the brain has been associated with aberrant neuronal network activity and poor cognitive performance in preclinical mouse models of Alzheimer’s disease (AD). Presently, our understanding of the mechanisms driving pathology-associated neuronal dysfunction and impaired information processing in the brain remains incomplete. Here, we assessed the impact of advanced Aβ pathology on spatial information processing in the medial entorhinal cortex (MEC) of 18-month *App ^NL-G-F/NL-G-F^* knock-in (APP KI) mice as they explored contextually novel and familiar open field arenas in a two-day, four-session recording paradigm. We tracked single unit firing activity across all sessions and found that spatial information scores were decreased in MEC neurons from APP KI mice versus those in age-matched C57BL/6J controls. MEC single unit spatial representations were also impacted in APP KI mice. Border cell firing preferences were unstable across sessions and spatial periodicity in putative grid cells was disrupted. In contrast, MEC border cells and grid cells in Control mice were intact and stable across sessions. We then quantified the stability of MEC spatial maps across sessions by utilizing a metric based on the Earth Mover’s Distance (EMD). We found evidence for increased instability in spatially-tuned APP KI MEC neurons versus Controls when mice were re-exposed to familiar environments and exposed to a novel environment. Additionally, spatial decoding analysis of MEC single units revealed deficits in position and speed coding in APP KI mice in all session comparisons. Finally, MEC single unit analysis revealed a mild hyperactive phenotype in APP KI mice that appeared to be driven by narrow-spiking units (putative interneurons). These findings tie Aβ-associated dysregulation in neuronal firing to disruptions in spatial information processing that may underlie certain cognitive deficits associated with AD.

## Introduction

The entorhinal cortex (EC) and hippocampus play pivotal roles in the processing and integration of spatial information within the mammalian brain (O’Keefe and Nadel, 1978; Witter and Moser, 2006; Igarashi, 2016; Stensola and Moser, 2016). Select neurons in these regions, i.e., grid, border, head-direction, speed, and place cells, are spatially tuned and exhibit distinctive firing characteristics during active exploration. Collectively, these cells are thought to contribute to the formation of a cognitive map of space through the utilization of both spatial and temporal coding schemes (Robinson et al., 2017; Behrens et al., 2018; Poulter et al., 2018). Notably, the EC is also a principal site of aggregated pathology and neuronal loss observed in Alzheimer’s disease (AD), a debilitating neurodegenerative disease characterized symptomatically by a progressive and irreversible decline in cognitive function (Braak and Braak, 1991; Van Hoesen et al., 1991; Braak and Braak, 1995; Gomez-Isla et al., 1996). Two pathological hallmarks of AD in the brain include the accumulation of hyperphosphorylated, misfolded tau proteins into neurofibrillary tangles (NFT), along with the deposition of amyloid beta (Aβ) into extracellular plaques (Gallardo and Holtzman, 2019; Long and Holtzman, 2019). Alterations in navigational strategies and reduced grid-cell-like representations in EC have been reported in young adults at genetic risk for AD (*APOE-*ε*4* carriers) and in healthy, aged human subjects (Kunz et al., 2015; Stangl et al., 2018). It is possible then that early pathological conditions in the course of AD progression may compromise spatial coding mechanisms within the EC and hippocampus, ultimately leading to deficits in path integration and increased spatial disorientation (Parizkova et al., 2018; Segen et al., 2022; Igarashi, 2023).

Preclinical studies utilizing *in vivo* single unit recordings in rodent models of AD pathology have yielded important insights on spatial cell dysfunction and their possible contributions to spatial learning and memory deficits (Silva and Martinez, 2022). For instance, hexagonal grid-like representations in medial entorhinal cortex (MEC) grid cells were shown to be severely disturbed in mouse models of Aβ pathology (Jun et al., 2020; Ying et al., 2022; Ying et al., 2023) and tau pathology (Fu et al., 2017; Ridler et al., 2020). In the hippocampus, Aβ and tau pathology are associated with reduced spatial coding in place cells, as well as impaired remapping of place fields in mice exposed to changes in environmental context (Cacucci et al., 2008; Zhao et al., 2014; Mably et al., 2017; Jun et al., 2020). While these findings have contributed valuable information on pathology-associated spatial cell dysfunction, spatial cell types other than place and grid cells are likely impacted by pathology as well (*e.g.,* border, speed, head-direction cells, *etc.*) (Ying et al., 2022; Ying et al., 2023), along with nonperiodic spatial cells (Zhang et al., 2013; Diehl et al., 2017; Miao et al., 2017) and non-spatial cells that contribute to spatial coding (Hangya et al., 2010; Buetfering et al., 2014; Miao et al., 2017). Aβ and tau pathology may also impact the stability of spatial information coding during exploration of novel and/or familiar environments. It is therefore important to investigate 1.) the quality of spatial coding features in mice exhibiting AD pathology and 2.) the stability of spatial coding in these mice over time, regardless of information quality, to better understand circuit mechanisms underlying cognitive impairment.

Functional magnetic resonance imaging (fMRI) studies have shown that individuals at risk for AD (e.g. amnestic mild cognitive impairment (aMCI), *APOE-*ε*4* carriers) (Bondi et al., 2005; Dickerson et al., 2005; Sperling et al., 2010; Putcha et al., 2011; Bakker et al., 2012), as well as pre-symptomatic carriers of familial AD (FAD) mutations (Quiroz et al., 2010), exhibit pronounced hyperactivity in the hippocampus during task-related encoding. Coupled with an increased prevalence of unprovoked seizures in AD patients and a faster rate of cognitive decline (Friedman et al., 2012; Vossel et al., 2013; DiFrancesco et al., 2017; Vossel et al., 2017), overexcitation in large networks of neurons may be a clinically relevant feature of AD (Busche and Konnerth, 2015; Zott et al., 2018). Preclinical studies in rodent models of AD pathology have shown that increased neuronal activity plays a key role in both Aβ accumulation (Cirrito et al., 2005; Yamamoto et al., 2015; Yuan and Grutzendler, 2016) and accelerated tau pathology in vivo (Yamada et al., 2014; Wu et al., 2016; Schultz et al., 2018). Moreover, increased seizure susceptibility, neuronal hyperactivity and network dysfunction have been reported in several rodent models of Aβ pathology (Palop et al., 2007; Busche et al., 2008; Minkeviciene et al., 2009; Busche et al., 2012; Verret et al., 2012; Petrache et al., 2019; Hijazi et al., 2020; Johnson et al., 2020; Rodriguez et al., 2020; Sosulina et al., 2021; Inayat et al., 2023; Soula et al., 2023; Rajani et al., 2024). Aβ-associated hyperactivity may indirectly impact spatial coding in the EC and hippocampus by creating a dysfunctional network environment, whereby spatial and non-spatial cells are affected. Alternatively, elevated levels of Aβ pathology (e.g. soluble Aβ species) may directly interact with spatial cells in these regions, disturbing cell firing patterns and distorting spatial tuning properties.

While much is known about the pathophysiology of Aβ and tau progression in the brain, a comprehensive understanding of the mechanisms underlying pathology-associated dysfunction in spatial coding remains incomplete. In this study, we show that advanced Aβ pathology in the brain is associated with impaired spatial information coding in 18-month *App ^NL-G-F/NL-G-F^* knock-in (APP KI) mice during exploration of novel and familiar contextual environments. Specifically, we show that MEC neurons in aged APP KI mice exhibit decreased spatial information content and impaired firing stability in our paradigm, in addition to poor position and speed coding relative to neurons in age-matched Control mice. We also found a mild hyperactivity phenotype in the MEC of APP KI mice. Altogether, these results provide evidence that advanced Aβ pathology in aged mice disturbs spatial information coding in MEC during exploration of novel and familiar environments.

## Results

### MEC neurons in aged APP KI mice exhibit poor spatial information content in familiar and novel contexts

APP KI mice express humanized mutant amyloid precursor protein (APP) at physiologically relevant levels, leading to aggressive age-associated amyloidosis and Aβ plaque formation in the brain beginning as early as 3-4 months (Saito et al., 2014; Benitez et al., 2021). APP KI mice with significant Aβ pathology (6-13 months) exhibit deficits in context discrimination and spatial learning and memory, as well as impaired remapping in CA1 place cells and MEC grid cells (Saito et al., 2014; Mehla et al., 2019; Johnson et al., 2020; Jun et al., 2020; Inayat et al., 2023). These findings suggest an association between Aβ pathology, spatial cell dysfunction, and cognitive impairment. However, little is known about general spatial information quality in these mice and whether the spatial information code is stable over long periods of time and in familiar versus novel environments.

To assess the impact of advanced Aβ pathology on stability of spatial information coding over large timescales, we performed in vivo MEC recordings in 18-month APP KI mice and age-matched C57BL/6J (control) mice as they explored square open field arenas in a sequence of sessions called recording blocks (**Figure 1A-B**). Recording blocks were comprised of four individual recording sessions spread over two consecutive days, with two sessions performed each day. Sessions 1 and 2 were recorded on Day 1 in the morning and afternoon, respectively. Sessions 3 and 4 were recorded in the morning and afternoon of the following day (Day 2). The intersession interval (ISI) was 4-6hr for same-day recordings (S1 vs S2, S3 vs S4) and 16-18hr for S2 vs S3. Importantly, the contextual features of the open field arena were kept constant for S1 through S3, then changed for S4 to promote remapping events. All arena contexts used in these experiments are shown in **Supplemental Figure 1**.

**Figure 1.**
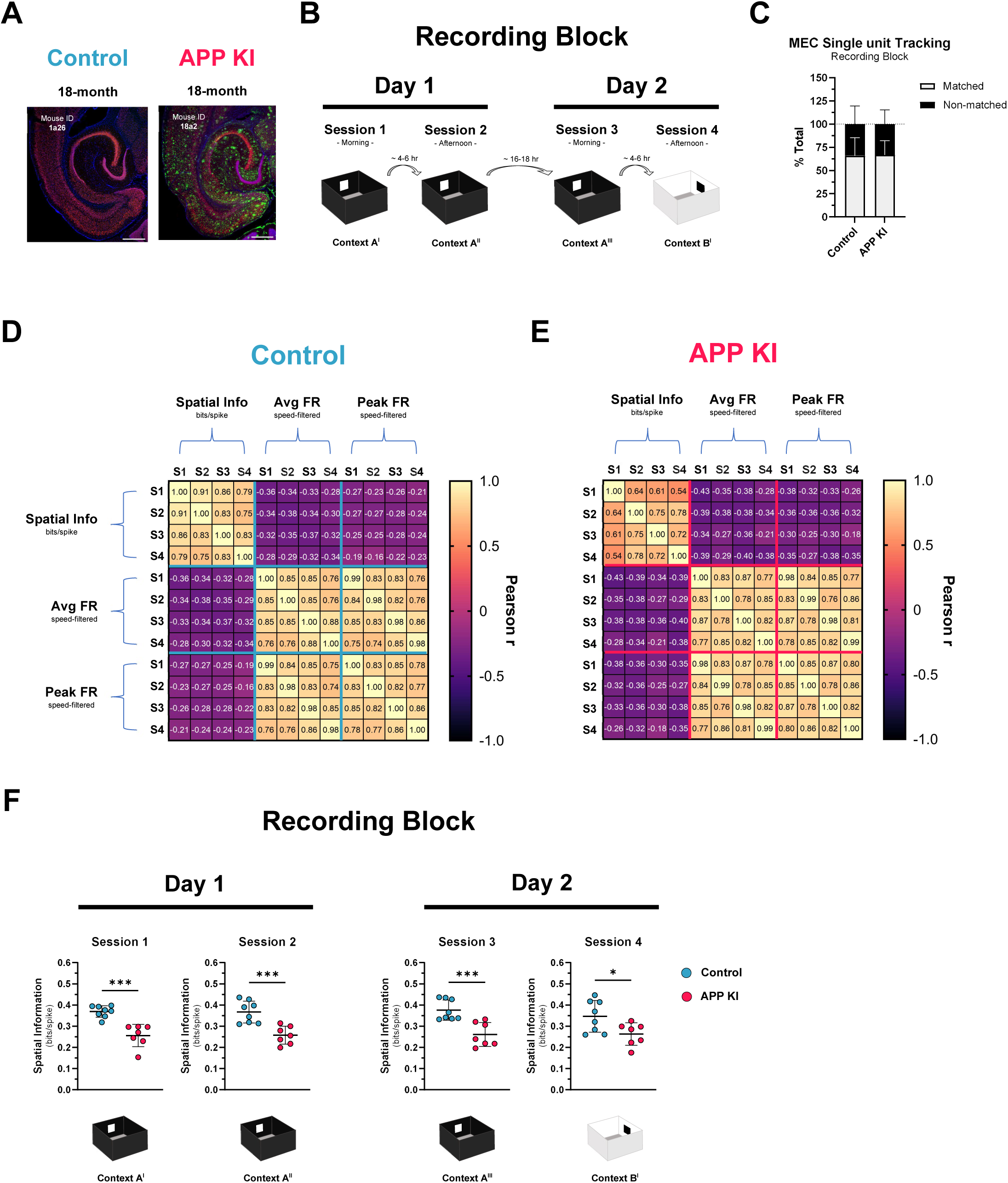
Aged *APP ^NL-G-F/NL-G-F^*mice exhibit impaired spatial information processing in the medial entorhinal cortex. ***A*.** Immunofluorescence staining in horizontal brain sections of 18-month *App ^NL-G-F/NL-G-F^* knock-in (APP KI) mice show significant amyloid beta (Aβ) pathology throughout the entorhinal cortex and hippocampus. Age-matched, non-transgenic C57BL/6J control mice did not exhibit any amyloid accumulation. Mouse IDs appear at upper left of image panels. Aβ, green; NeuN, red; Hoechst, blue. Scale bar, 500µm. ***B*.** Experimental design to assess stability of spatial coding in novel and familiar contextual environments. 18-month APP KI mice and age-matched Control mice were implanted with electrodes in the medial entorhinal cortex (MEC) and allowed to freely explore an open field arena while single unit and local field potential (LFP) data were collected. The paradigm consisted of recording activity in four recording sessions across two days. Session 1, Context A^I^, Novel; Session 2, Context A^II^, Familiar; Session 3, Context A^III^, Familiar; Session 4, Context B^I^, Novel. Superscript indicates number of exposures to the context in a given recording block. All square arenas measure 50cm x 50cm. ***C*.** Individual single units collected from 18-month APP KI (Total, n=7 mice) and Control (Total, n=8 mice) were carefully matched across recording sessions by examining average waveforms and cluster positions in 2D feature space. ‘Matched’ single units were those units that were readily identified and consistent across all four recording sessions in a recording block. ‘Non-matched’ single units were defined as units that were readily identified in one or several sessions, but not all four sessions in a recording block. A total of 352 units were isolated in our recording block (Control, n=183 units; APP KI, n=169 units). A total of 226 units were classified as matched (Control, n=115 units; APP KI, n=111 units), and 126 were classified as non-matched (Control, n=68 units; APP KI, n=58 units). Percentages of matched and non-matched units per mouse were calculated and the group means were plotted. Gray bars, matched units. Black bars, non-matched units. No genotype differences were found in the % of matched or non-matched units tracked in our experiment. Unpaired t-test: *t*(13) = 0.0898, *P* = 0.9298. ***D-E*.** Correlation matrices were used to evaluate the cross-session relationships between speed-filtered average firing rates (AVG FR), peak firing rates, and spatial information (SI) scores in Control mice and APP KI mice. ***F*.** Mean SI scores were calculated per mouse for each session and group means were compared per session. MEC units recorded from APP KI mice exhibited decreased SI scores versus Control MEC units in Sessions 1, 2 and 3. Unpaired *t*-tests: Session 1, *t*(13) = 5.370, *p* < 0.001; Session 2, *t*(13) = 4.423, *p* < 0.001; Session 3, *t*(13) = 4.259, *p* < 0.001; Session 4, *t*(13) = 2.452, *p* < 0.05. Bar graphs represent mean ± standard deviation (SD). Scatter plots represent mean ± SD. * *p* < 0.05; ** *p* < 0.01; *** *p* < 0.001.

Individual MEC neurons were tracked across sessions in our recording blocks and classified as ‘matched’ if they reliably appeared in each session of the block. ‘Non-matched’ neurons were any units that did not appear in all four sessions of the block or those that could not be reliably matched. In total, 226 matched MEC neurons were recorded and analyzed for this study (Control, n=115 MEC single units; APP KI, n=111 MEC single units). A total of 126 MEC neurons could not be reliably matched across recording sessions and were not included in spatial coding analysis. We found no significant differences in the number of matched single units tracked (*p*>0.05) or non-matched single units tracked (*p*>0.05) between groups (**Figure 1C**). To examine task-relevant neuronal firing rates during active exploration, we applied a minimum-maximum speed filter (3-100 cm/sec) to the data to remove spiking activity during bouts of immobility (<3 cm/sec) and spurious jumps in activity (>100cm/sec). We then examined the relationship between speed-filtered average firing rates (AVG FRs), peak firing rates (PK FRs) and spatial information (SI) scores of matched MEC neurons in our dataset (**Figure 1D-E**). SI scores were calculated using Skaggs’ formula and reflect the extent to which a recorded unit’s firing can predict the location of the animal in the arena (Skaggs et al., 1993; Skaggs et al., 1996). First, correlation matrices revealed strong negative relationships between firing rates and SI scores across sessions, regardless of genotype. No discernable differences in relationship strength were found between groups in MEC AVG FRs or PK FRs across recording sessions. However, MEC neurons from APP KI mice showed a weak relationship in SI scores across recording sessions compared to Control mice. Looking closer, we found significant group differences in SI scores, with APP KI mice exhibiting lower scores than Control mice in all four sessions (S1, *p*<0.001; S2, *p*<0.001; S3, *p*<0.001; S4, *p*<0.05) (**Figure 1F**). Cross-session correlations of individual MEC single unit SI scores were then performed to assess their stability through the recording block (**Supplemental Figure 2**). SI scores were highly correlated across sessions in Control mice (Pearson’s r means range: 0.84 - 0.92), whereas SI score cross-session correlations were relatively weaker in APP KI mice (Pearson’s r means range: 0.66 – 0.77). Group differences were detected in S1 vs S2 (*p*<0.05), and S1 vs S3 (*p*<0.05) comparisons (**Supplemental Figure 2A-D**), with no differences detected in all other session comparisons.

Speed-filtered (3-100 cm/sec) MEC oscillatory activity was also analyzed in our mice across recording sessions and compared between genotypes. However, no group differences were detected in percentage power of filtered LFP signals in the delta (1-3 Hz; *p*>0.05), theta (4-12 Hz; *p*>0.05), beta (13-20 Hz; *p*>0.05), low gamma (35-55 Hz; *p*>0.05), or high gamma (65-120 Hz; *p*>0.05) frequency ranges in these experiments (**Supplemental Figure 3**). To ensure gross locomotor activity did not differ between groups and in individual animals across our two-day recording block, we analyzed the position data from our mice as they freely explored the open field arenas during recording sessions (**Supplemental Figure 4**). We found no group differences in the total distance traveled (m), the percentage of arena coverage (%), or the average speed (cm/sec) of the mice across sessions in our recording block (*p*>0.05) (**Supplemental Figure 5A-D**). However, correlation matrices examining the relationships between these measures by session did reveal genotype differences in arena coverage vs total distance traveled and average speed (**Supplemental Figure 5E**). Those differences were explained by increased exploration of the arena center in APP KI mice versus Control mice (Zone x Genotype; *p*<0.01), with *post hoc* analyses revealing significant group differences in S1 (*p*<0.01) and S3 (*p*<0.05) (**Supplemental Figure 5F**). This effect was also reflected by the decreased exploration of arena boundaries in S1 (*p*<0.01) and S3 (*p*<0.05).

Together, these data demonstrate that MEC neurons in 18-month APP KI mice exhibit relatively poor spatial information during exploration of a novel environment (S1, Context A^I^), and that information quality does not improve with repeated exposures to the same environment over time (S2, Context A^II^; S3, Context A^III^), or upon exposure to an additional novel environment (S4, Context B^I^). We also found that informational stability in APP KI MEC neurons appeared to be compromised early in this recording paradigm, as shown by cross-session correlations of SI scores. These effects could not be attributed to gross differences in locomotor activity, though aged APP KI mice did spend more time exploring the center of the arenas versus Control mice.

### MEC spatial representations are unstable across familiar and novel contexts in aged APP KI mice

To examine the quality and stability of spatial representations of individual MEC neurons across sessions in our recording paradigm, we identified matched single units that exhibited grid, border, and/or head-directional (HD) tuning in at least one recording session of several recording blocks available per mouse (**Figure 2**). MEC grid cells in 18-month Control mice exhibited hexagonal firing patterns with well-formed firing fields that appeared stable across all recording sessions (**Figure 2A**) (Fyhn et al., 2004; Hafting et al., 2005). In contrast, grid cell tuning was significantly disturbed in 18-month APP KI mice, supporting previous findings in relatively younger knock-in animals (Jun et al., 2020). These nonperiodic, spatially-tuned APP KI MEC single units displayed scattered firing patterns in the arenas, with dysmorphic firing fields that were unstable across all recording sessions (**Figure 2B**). MEC border cells were identified by the appearance of a strong firing field proximal to an environmental boundary (Solstad et al., 2008). Control MEC border cells displayed well-defined firing fields that were stable across sessions (**Figure 2C**). Interestingly, border cell firing was also present in APP KI mice, though border preference was largely unstable across sessions. Examples of border cell instability in APP KI mice include the rotation and broadening of firing fields and the appearance or disappearance of spatial tuning during exploration of both familiar and novel arena contexts (**Figure 2D**). Notably, we did not detect group differences in tuning properties of MEC HD cells between APP KI and Control mice. Single units were classified as HD cells if they preferentially displayed tuning toward a particular direction in polar plots (Taube et al., 1990; Sargolini et al., 2006). HD cells were strongly tuned and stable across all recording sessions in both APP KI and Control mice (**Figure 2E-F**).

**Figure 2.**
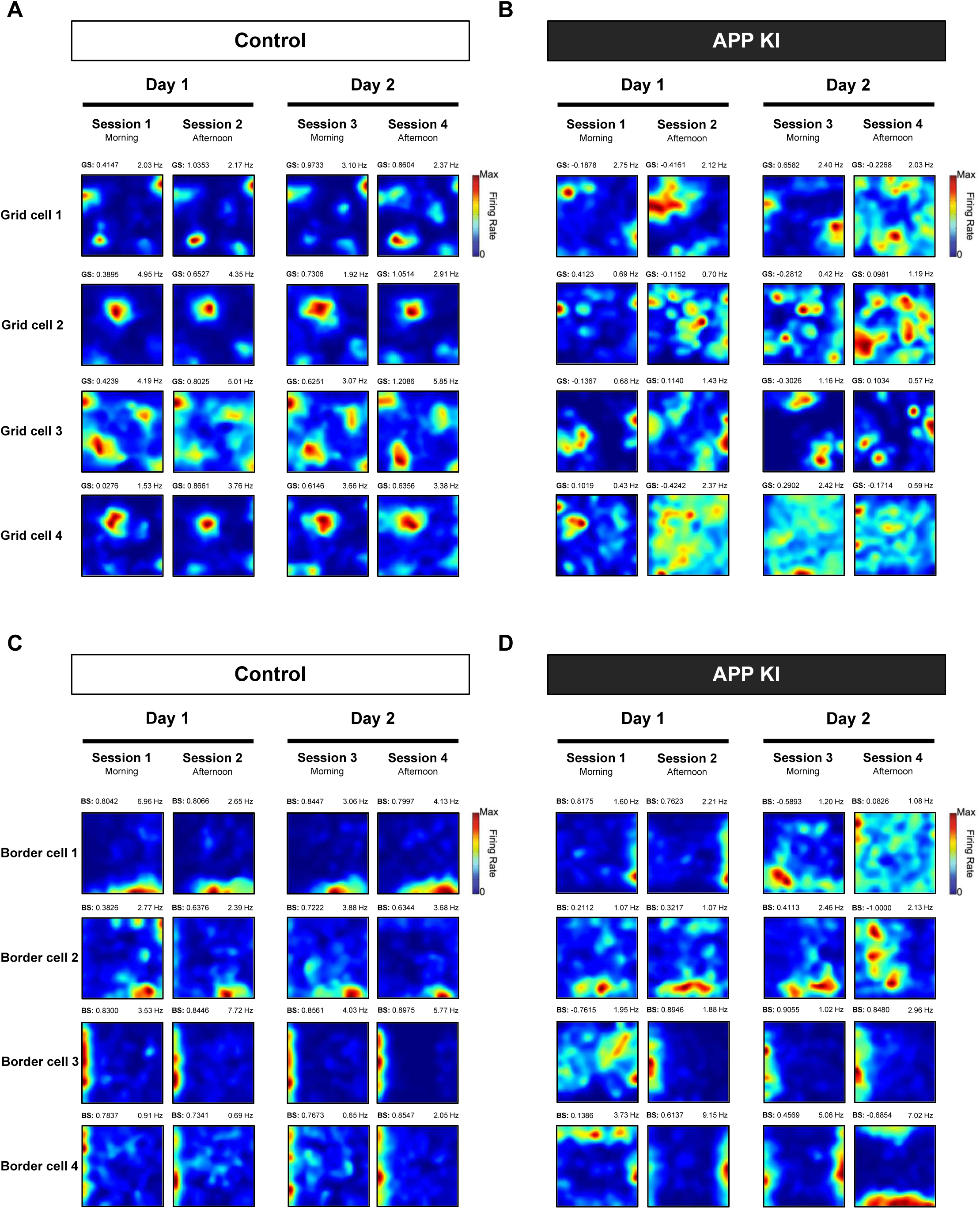
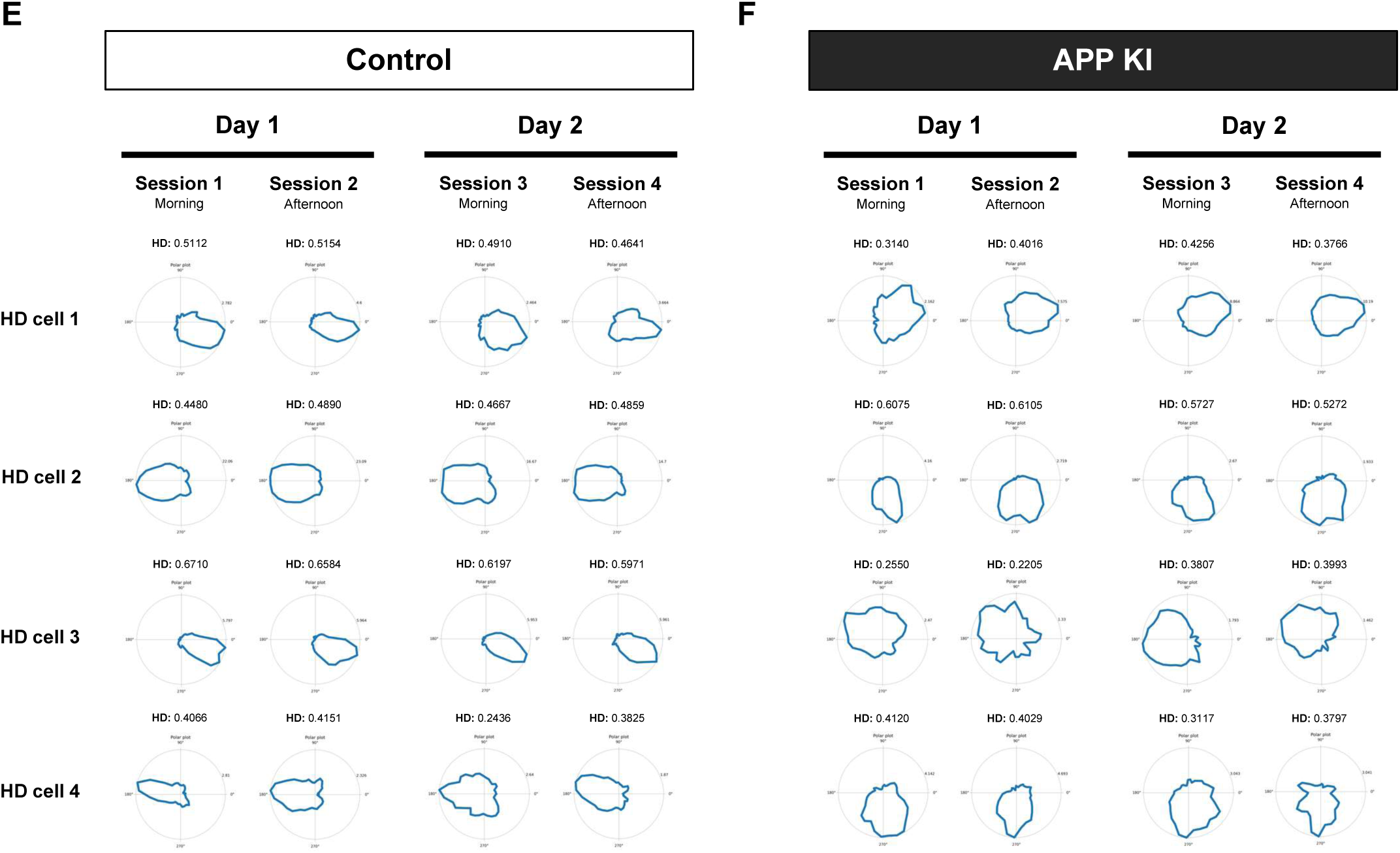
Aged *App ^NL-G-F/NL-G-F^* mice exhibit impaired stability of spatial tuning in MEC grid cells and border cells, but not head-direction cells. Individual MEC neurons exhibiting spatial tuning features were identified and tracked across sessions in several recording blocks. These neurons were classified as grid, border or head-direction cells. ***A-B***. Firing rate maps of representative MEC grid cells are shown for 18-month Control mice and age-matched APP KI mice. Grid-like representations appeared intact and stable in aged Control mice across the two-day recording sessions, while firing patterns appeared scattered and unstable in APP KI mice. Grid scores (GS) appear at the top left of the rate maps, while speed-filtered (3-100 cm/sec) peak firing rates appear at top right. Animal IDs, recording block IDs, and cell IDs are as follows: Control: 1-24, S12-13, T2C1. 1-25, S17-18, T4C1. 1-28, S14-15, T2C1. 1-25, S21-22, T4C1. APP KI: 1-13, S34-35, T3C4. 1a27, S14-15, T4C1. 1-13, S33-34, T3C6. 1a37, S22-23, T3C7. ***C-D*.** Firing rate maps of representative MEC border cells are shown for aged Control and APP KI mice. Geometric border-like representations appeared intact and stable in Control mice across the two-day recording sessions. In aged APP KI mice, border cell firing appeared intact but with broader and unstable tuning compared to border cells in Controls. Border scores (BS) appear at the top left of the rate maps, while speed-filtered peak firing rates appear at top right. Animal IDs, recording block IDs, and cell IDs are as follows: Control: 1-25, S29-30, T4C4. 1-28, S14-15, T4C1. 1-34, S12-13, T2C1. 1-28, S14-15, T4C2. APP KI: 1-29, S11-12, T4C1. 1a33, S17-18, T3C1. 1a33, S9-10, T4C1. 1a27, S32-33, T3C2. ***E-F*.** Head-directional tuning in MEC neurons appeared intact and stable across recording sessions in both aged Control and APP KI mice. Head-direction (HD) scores appear at top of the polar plots. Animal IDs, recording block IDs, and cell IDs are as follows: Control: 1a21, S6-7, T2C2. 1a21, S14-15, T2C2. 1a21, S14-15, T2C1.1-28, S21-22, T4C2. APP KI: 1-13, S17-18, T3C2. 1-13, S26-27, T2C1. 1-15, S38-39, T3C1. 1-14, S38-39, T3C1.

We then asked whether representational stability could be measured in all single units collected in our recording block, not just in conventional MEC spatial cell types. To quantify the stability of spatial representations in MEC neurons across sessions in our recording block, we calculated ratemap differences in key session comparisons using a noise-resistant, rate-robust metric based on the Earth Mover’s Distance (EMD) (**Figure 3**) (Aoun et al., 2023). The EMD is a measure from optimal transport theory that reflects the minimal cost required to transform one distribution into another, where the cost is proportional to the amount of “earth” moved and the distance it is moved (Bonneel et al., 2015). In this context, the ratemap firing densities in our recording sessions represent “earth” and the distance moved between two ratemaps being compared represents the change in that density. Here, we use the ‘whole map EMD’ as a measure of dissimilarity between ratemaps consisting of binned unit spikes where the arena dimensions were fixed (32×32) across animals and arena contexts (**Figure 3A**).

**Figure 3.**
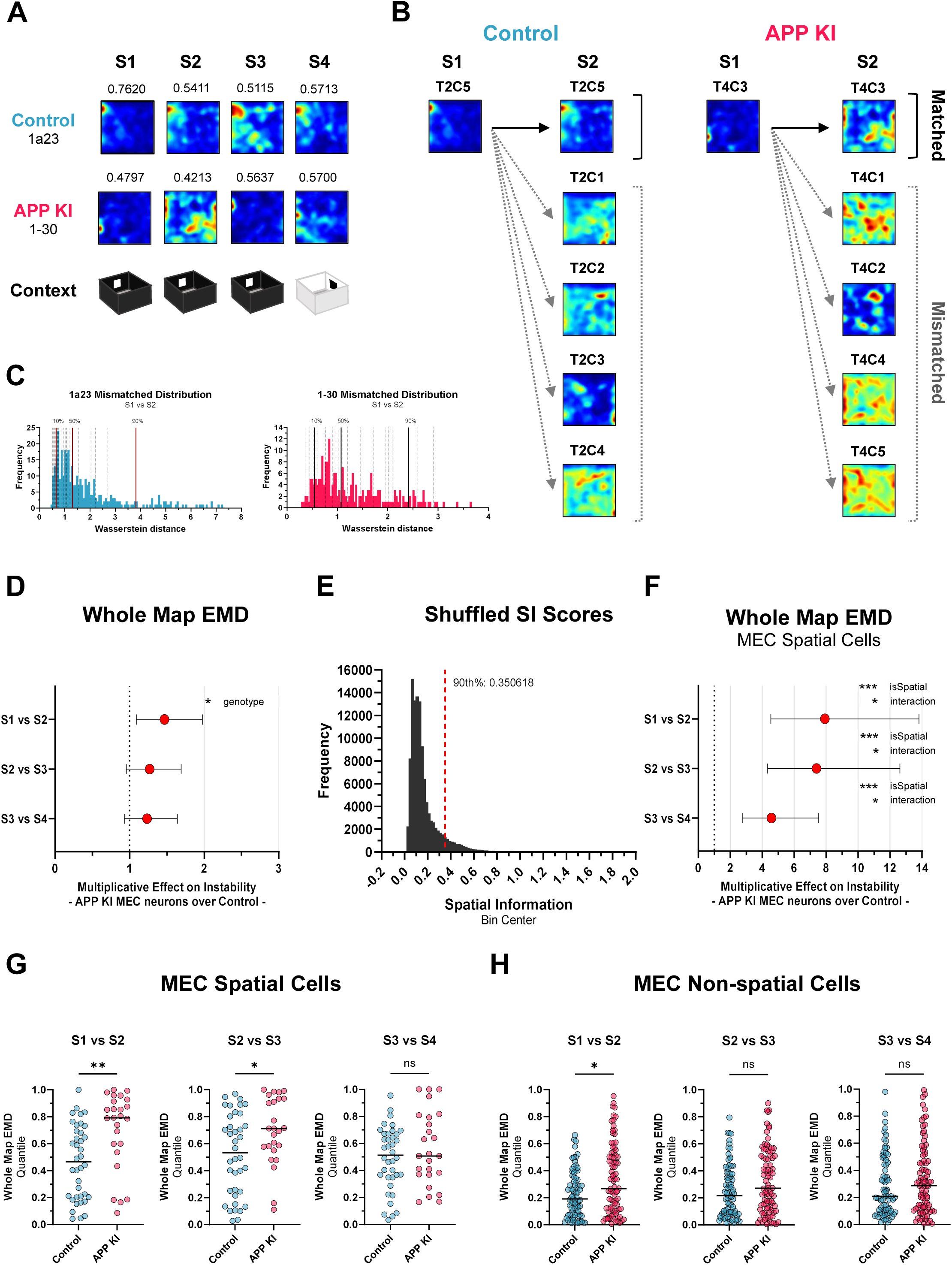
MEC spatial cells in aged APP KI mice exhibit increased representational instability in familiar environments. The spatial dissimilarity between single unit ratemaps in our recording block was quantified using the Earth Mover’s Distance (EMD). Here, the goal is to find the minimum cost required to transform one distribution into another (session comparison). In our recording block, a single unit’s binned firing density is transformed into a distribution representing ‘earth’ and the ‘distance’ represents the change in that distribution. ***A***. All four recording session ratemaps from two mice in the recording block are shown. *Top*, Control, 1a23. *Bottom*, APP KI, 1-30. Spatial information (SI) scores for each session are shown atop each map. ***B-C*.** Whole map EMD scores (Wasserstein distances) were approximated using the sliced EMD and then converted into quantiles for each animal based on an EMD distribution of mismatched single units. For reference, frequency distribution plots are shown for mice 1a23 (*left*, blue) and 1-30 (*right*, red) in the S1 vs S2 comparison. The quantile values for matched MEC units appear as hashed vertical lines in the plots. Solid red vertical lines represent the 10^th^, 50^th^ and 90^th^ percentile of quantile values from mismatched units in the recording session comparison. ***D*.** A mixed-effects beta regression model was used to determine variable effects on the whole map EMD. Genotype served as a fixed effect with random effects adjusted for the subject (Animal ID) and session comparison. The resulting odds ratio plot reflects the multiplicative effect on instability of quantiles representing APP KI MEC neurons over Controls. We found an effect of genotype on the S1 vs S2 comparison (OR=1.467, *p*=0.011), but no differences on S2 vs S3 (OR=1.269, *p*>0.05) or S3 vs S4 (OR=1.234, *p*>0.05) comparisons. ***E*.** Spike times were misaligned from *x,y* positions for all matched MEC units in Session 1 of our recording block. Each unit was shuffled 1,000X times and a frequency distribution of circularly shuffled SI scores was plotted. Units with true SI scores above the 90^th^ percentile (0.3506) of this distribution were classified as spatial cells, while those that fell below were classified as non-spatial cells (Control: n=38 spatial units, n= 77 non-spatial units; APP KI: n=25 spatial units, n=86 non-spatial units. ***F*.** Introduction of an additional variable into the model labelling single unit session comparisons as spatial or non-spatial (isSpatial) yielded the following results. Significant interaction effects of genotype and isSpatial variables were detected for each session comparison (S1 vs S2, OR=7.92, *p*<0.05; S2 vs S3, OR=7.40, *p*<0.05; S3 vs S4, OR=4.58, *p*<0.05). The isSpatial variable had a significant main effect on all session comparisons (S1 vs S2, *p*<0.001; S2 vs S3, *p*<0.001; S3 vs S4, *p*<0.001), while genotype did not (S1 vs S2, *p*>0.05; S2 vs S3, *p*>0.05; S3 vs S4, *p*>0.05). ***G*.** Whole map EMD quantile values for MEC spatial cells were compared between genotype for each session comparison using two-tailed Mann-Whitney *U* tests. Quantile values were significantly larger in APP KI MEC spatial cells versus Control spatial cells in S1 vs S2 (*U*=245.5, *p*<0.01) and S2 vs S3 (*U*=305, *p*<0.05), but not in S3 vs S4 (*U*=407, *p*>0.05) comparisons. ***H*.** Quantile values of MEC non-spatial cells were also compared across genotype. Whole map EMD quantile values were significantly larger in APP KI MEC non-spatial cells versus Control non-spatial cells on S1 vs S2 (*U*=2,588, *p*<0.05) only. No genotype difference in quantile values were detected on S2 vs S3 (*U*=2,792, *p*>0.05) or S3 vs S4 (*U*=2,987, *p*>0.05) comparisons. For EMD analysis, ratemap bins were fixed across all animals and sessions (32X32 bins). EMD, Earth Mover’s Distance. SI, spatial information. OR, Odds Ratio. Red datapoints in EMD OR plots indicate OR values with full range confidence intervals. Scatter plots represent median with full data range. * *p*<0.05; ** *p*<0.01. *** *p*<0.001; ns, not significant.

Since raw EMD scores vary depending on both representational stability and intrinsic properties of the neurons under consideration, we first standardized the EMD scores by converting them into quantiles for each animal based on reference EMD distributions of mismatched units per session comparison (origin session vs target session) (**Figure 3B-C**). We then ran a mixed-effects beta regression model for the whole map EMD using the quantiles for each session comparison. In computing the odds ratio for the whole map EMD, we found evidence for increased instability in APP KI MEC neurons versus Controls on the first re-exposure to Context A in the recording sequence (S1 vs S2: Odds Ratio = 1.47, *p*<0.05), but not on the second re-exposure to Context A the following day (S2 vs S3: Odds Ratio = 1.27, *p*>0.05) or on exposure to a novel Context B (S3 vs S4: Odds Ratio = 1.23, *p*>0.05) (**Figure 3D**, **Table 1A**). However, when we introduced a variable into the model that identified matched MEC single units as ‘spatial’ versus ‘non-spatial’ using a circular shuffling method (**Figure 3E**), significant interaction effects were detected for each session comparison (S1 vs S2: *p*<0.05; S2 vs S3: *p*<0.05; S3 vs S24: *p*<0.05) (**Figure 3F**, **Table 1B**). MEC spatial units in APP KI mice that exceeded the 90^th^ percentile of circularly shuffled SI scores were ∼7.5X more likely than Control spatial units to exhibit representational instability during re-exposure to a familiar environment (S1 vs S2: Odds Ratio = 7.92; S2 vs S3: Odds Ratio = 7.40), and were ∼4.5X more likely than Control units to exhibit instability during exposure to a novel environment (S3 vs S4: Odds Ratio = 4.60) in the recording paradigm. The whole map EMD quantile values for MEC spatial cells were then compared between genotype for each session comparison (**Figure 3G**). Quantile values were significantly larger in APP KI MEC spatial cells versus Control spatial cells in S1 vs S2 (*p*<0.01) and S2 vs S3 (*p*<0.05) comparisons, but not in S3 vs S4 (*p*>0.05). When examining the non-spatial cell population, significant genotype differences in quantile values were also found in S1 vs S2 (*p*<0.05), but not in S2 vs S3 or S3 vs S4 comparisons (*p*>0.05) (**Figure 3H**).

**Table 1.**
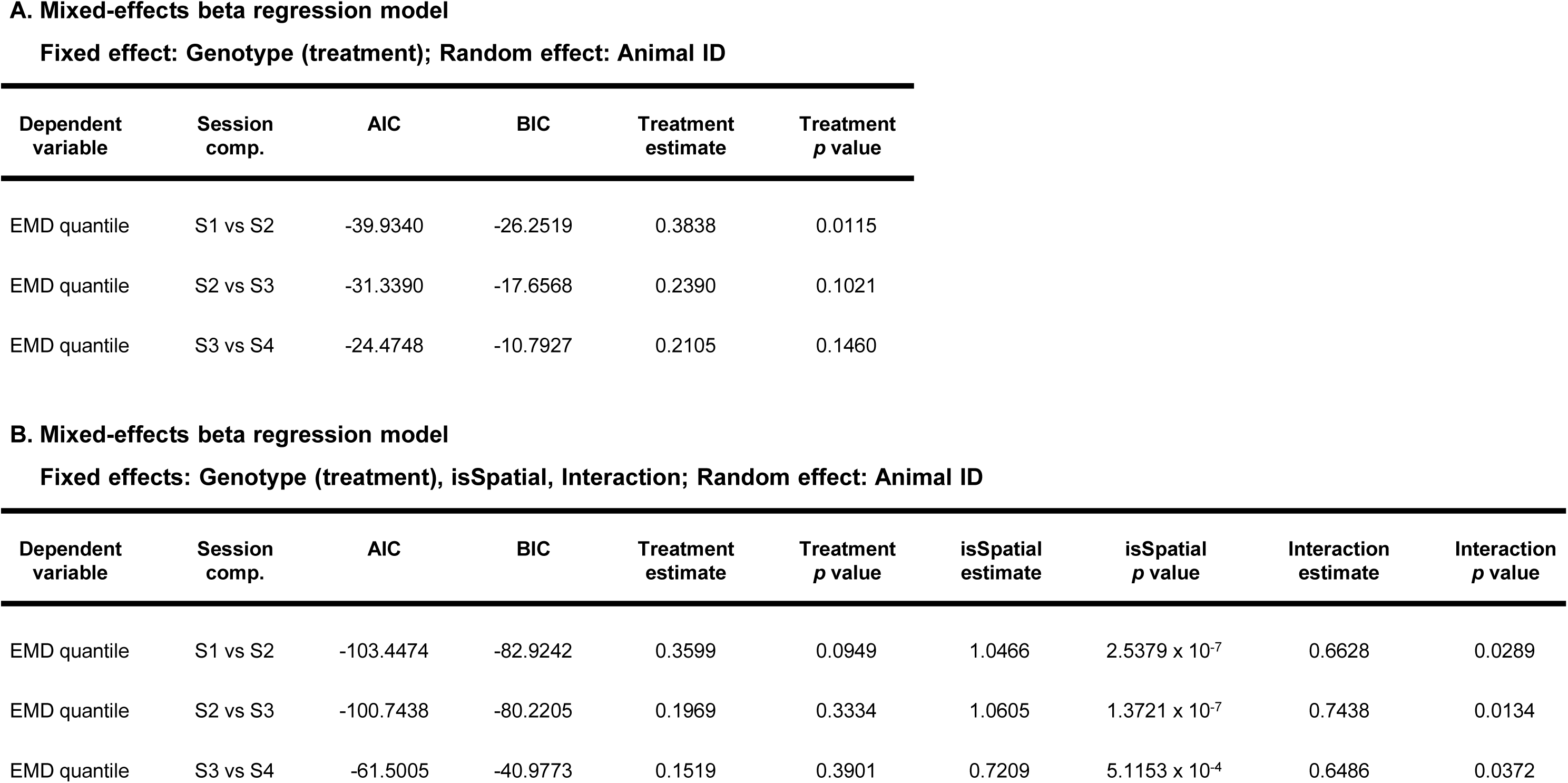
Summary statistics for mixed-effects beta regression models. The table shows the genotype (treatment), isSpatial, and interaction estimates and associated *p* values for each session comparison in Figure 3. AIC and BIC values are also shown. ***A*.** Values correspond to Figure 3D. ***B*.** Values correspond to Figure 3F. AIC, Akaike Information Criteria. BIC, Bayesian Information Criterion. EMD, Earth Mover’s Distance.

**Fixed effect: Genotype (treatment); Random effect: Animal ID**
| Dependent variable | Session comp. | AIC | BIC | Treatment estimate | Treatment *p* value |
| --- | --- | --- | --- | --- | --- |
| EMD quantile | S1 vs S2 | -39.9340 | -26.2519 | 0.3838 | 0.0115 |
| EMD quantile | S2 vs S3 | -31.3390 | -17.6568 | 0.2390 | 0.1021 |
| EMD quantile | S3 vs S4 | -24.4748 | -10.7927 | 0.2105 | 0.1460 |

**Table 1. Summary statistics for mixed-effects beta regression models.**
| Dependent variable | Session comp. | AIC | BIC | Treatment estimate | Treatment *p* value | isSpatial estimate | isSpatial *p* value | Interaction estimate | Interaction *p* value |
| --- | --- | --- | --- | --- | --- | --- | --- | --- | --- |
| EMD quantile | S1 vs S2 | -103.4474 | -82.9242 | 0.3599 | 0.0949 | 1.0466 | 2.5379 x 10<sup>-7</sup> | 0.6628 | 0.0289 |
| EMD quantile | S2 vs S3 | -100.7438 | -80.2205 | 0.1969 | 0.3334 | 1.0605 | 1.3721 x 10<sup>-7</sup> | 0.7438 | 0.0134 |
| EMD quantile | S3 vs S4 | -61.5005 | -40.9773 | 0.1519 | 0.3901 | 0.7209 | 5.1153 x 10<sup>-4</sup> | 0.6486 | 0.0372 |

An alternative form of shuffling that shifted firing fields within a ratemap was also applied to our MEC dataset to identify spatial units from a distribution of field shuffled SI scores (**Supplemental Figure 6A**) (Grieves et al., 2021). Again, we found significant interaction effects for each session comparison in our mixed-effects model (S1 vs S2: *p*<0.05; S2 vs S3:*p*<0.05; S3 vs S24:*p*<0.05), where MEC spatial units in APP KI mice exceeding the 90^th^ percentile of field shuffled SI scores were ∼8-9X more likely than Control units to exhibit representational instability in a familiar environment (S1 vs S2: Odds Ratio = 9.17; S2 vs S3: Odds Ratio = 7.78) (**Supplemental Figure 6B; Supplemental Table 1**). When exposed to a novel environment, APP KI spatial units were ∼5X more likely than Control units to exhibit ratemap instability (S3 vs S4: Odds Ratio = 5.18). Quantile values were significantly larger in field shuffle-identified APP KI MEC spatial cells versus Control in S1 vs S2 (*p*<0.001) and S2 vs S3 (*p*<0.05) comparisons, but not in S3 vs S4 (*p*>0.05) (**Supplemental Figure 6C**). Quantile values were also significantly larger in APP KI MEC non-spatial cells versus Control in S1 vs S2 (*p*<0.05), but not in S2 vs S3 or S3 vs S4 comparisons (*p*>0.05) (**Supplemental Figure 6D**).

In our recording paradigm, advanced Aβ pathology appeared to impact the spatial tuning properties of MEC spatial cells differently. Grid cells were predictably impaired, while border and HD cells showed strong tuning properties in all session recordings. However, border cells showed increased instability in both familiar and novel environments despite strong spatial tuning. HD tuning was strong and stable throughout the recording paradigm, regardless of genotype. When quantifying the representational stability of MEC spatial units in our dataset using the whole map EMD, we found that the effect of genotype on MEC neuron instability in familiar and novel contexts varied significantly depending on whether cells were classified as spatial versus non-spatial.

### MEC neurons in aged APP KI mice exhibit poor spatial coding for position and speed

Recent work has shown that MEC grid cells exhibit multiple firing fields during stationary treadmill running in rats, integrating information on distance run and elapsed time (Kraus et al., 2015). These data, among others, champion the notion that neurons in the EC and hippocampus are capable of responding to diverse combinations of task relevant variables (Pastalkova et al., 2008; MacDonald et al., 2011). This ability of neuronal ‘multitasking’ is termed mixed-selectivity (Fusi et al., 2016), a property theorized to ‘endow the computational horsepower needed for complex thought and action’. Mixed-selectivity can be probed by decoding neuronal activity at the population level, where contributions to information encoding are determined from activity patterns of large neuronal ensembles rather than individual neuron response properties (Stefanini et al., 2020).

Here, we asked if spatial information could be decoded from the activity patterns of many MEC neurons in our aged APP KI and Control mice as they explored familiar and novel contextual environments. To accomplish this, we used ensembles of one-versus-one linear support vector machines (SVMs) to classify MEC neural population vectors into 25 spatial bins (5×5). Spatial error was computed by imputing the center of each bin as the prediction and then calculating the mean squared error (MSE or L2) with respect to the imputed predictions. Pseudo-population decoding of animal head direction and speed information from the collective MEC activity was also performed using SVM ensembles. Speed error was computed as the absolute difference (L1 Loss) of bin centers to ground truth. These analyses yield three possibilities, which are depicted in **Figure 4A**: 1.) no coding: the neural patterns for different values of the analyzed variable (*e.g.*, different positions) are not reliably different from each other; hence they do not carry information about the analyzed variable; 2.) intact coding but no generalization across recording sessions: the neural patterns are reliably encoding information about the decoded variable, but they do so in different ways depending on the recording session (see how the linear classifier that separates orange vs. yellow points is different compared to the one separating purple vs. pink points in **Figure 4A**); 3.) intact decoding and generalization across sessions: different positions are reliably encoded in the neural patterns, and their encoding strategy is consistent across different sessions (the same linear classifier generalizes from orange vs. yellow to purple vs. pink). Generalization ability is assessed by testing the ability of the set of classifiers fitted in one session to decode the same variable in a second session (see Methods). In addition to the bin numbers chosen for illustration in **Figure 4A**, a parameter sweep of bin sizes was conducted. A graph of significance levels by session-comparison and bin number is included in the supplemental section.

**Figure 4.**
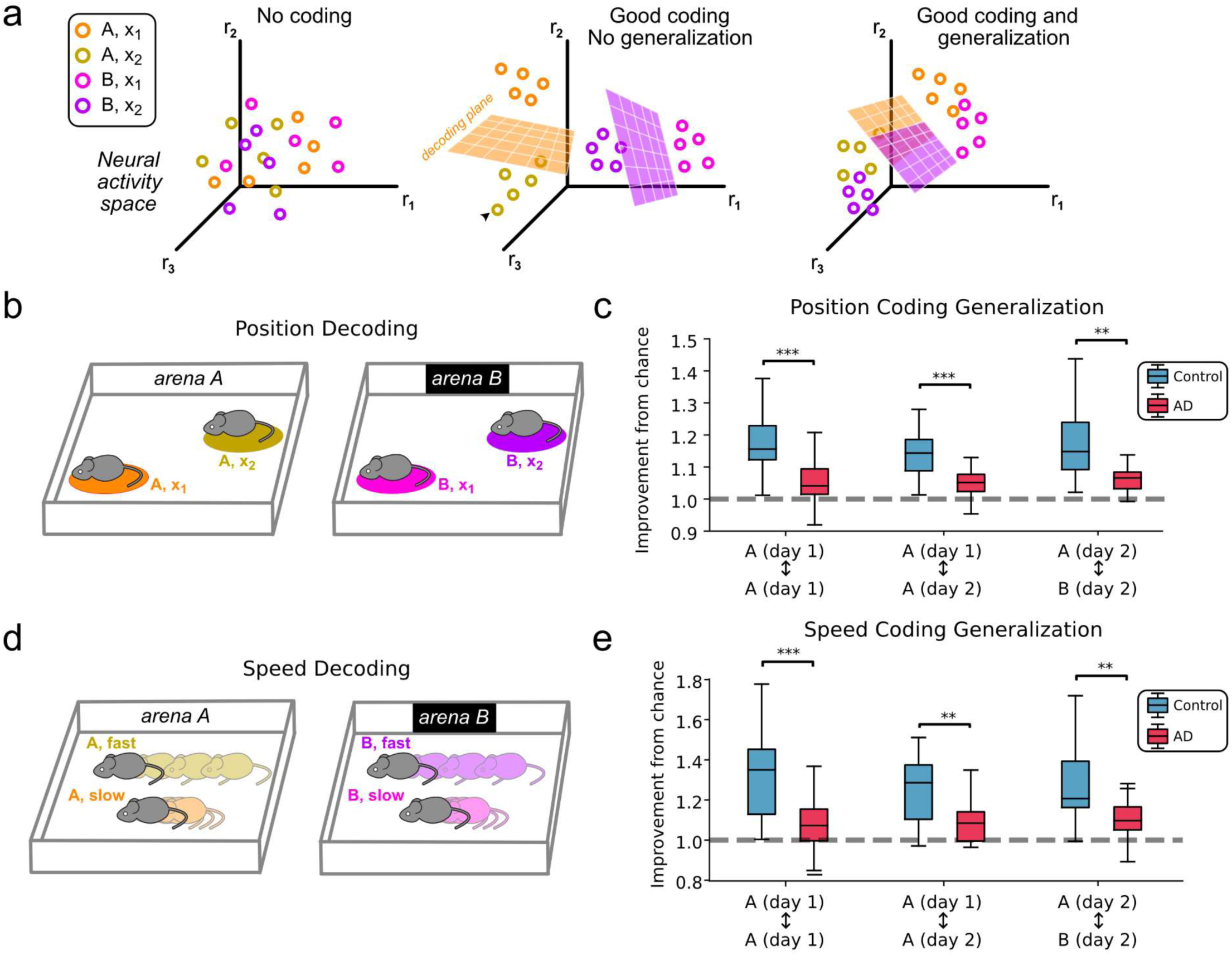
Spatial decoding is impaired across sessions in MEC neurons from aged *APP ^NL-G-F/NL-G-F^* mice. Linear support vector machines (SVMs) were used to decode discretized position and speed from a pooled set of neurons for 18-month-old animals in each group (APP KI vs Control). The SVMs were trained to decode data from within the same session (trained on the first half and assessed on the second half) and to decode across recording sessions (trained on the first session and assessed on the next session) ***A*.** Illustration of three possible decoding performance patterns for a given group. First, if no decoding is present, then the target categories will not be linearly separable. Second, if coding is possible, but the neural code does not generalize across sessions, then there will be a way to linearly separate the data by speed or position within each session, but the way of separating the data for each session will be different. Finally, if there is good coding and good generalization, then the way of separating the bins within each session will align with each other, allowing a model trained on one session to decode data from another session. ***B*.** Illustration of a mouse in different positions during two independent recording sessions. Note the change in arena context between sessions, depicted here as S3 (Context A) versus S4 (Context B). The colors correspond to the legend in A. ***C*.** Group differences in median decoding performance for position, expressed as an improvement from chance level, were detected in key session comparisons using Wilcoxon signed-rank tests (S1 vs S2; *p* < 0.001; S2 vs S3; *p* < 0.001; S3 vs S4; *p* < 0.01). ***D*.** Illustration depicting a mouse moving at different speeds during two independent recording sessions. The colors correspond to the legend in A. ***E*.** Group differences in median decoding performance for speed were detected in key session comparisons using Wilcoxon signed-rank tests (S1 vs S2; *p* < 0.001; S2 vs S3; *p* < 0.01; S3 vs S4; *p* < 0.01). Box plot and whiskers represent the interquartile range (IQR) and median improvement from chance values per genotype for each session comparison. ** *p* < 0.01; *** *p* < 0.001.

The decoding error ratios for MEC neurons from APP KI mice and Control mice were compared using Wilcoxon signed-rank tests across the decoding precision of different spatial positions or velocity bins. For generalization of position coding across sessions, we found significant group differences in the decoding error ratios on each session comparison (S1 vs S2, *p*<0.01; S2 vs S3, *p*<0.01; S3 vs S4, *p*<0.01), with Control MEC neurons showing greater improvement from chance versus APP KI MEC neurons (**Figure 4B-C**). A similar trend was found for generalized speed coding in our recording block, with Control MEC neurons outperforming APP KI MEC neurons on each session comparison (S1 vs S2, *p*<0.01; S2 vs S3, *p*<0.01; S3 vs S4, *p*<0.05) (**Figure 4D-E**). Independent runs of the decoding analyses were also performed on maps in which the arena space was divided into 16, 36 or 49 spatial bins (**Supplemental Figure 7**). In each bin condition, Control MEC neurons outperformed APP KI MEC neurons in position and speed decoding performance for all session comparisons.

To determine if poor cross-session generalization of MEC spatial coding in APP KI mice was due to deficits in information encoding within individual sessions, we halved each of the sessions in our recording block, trained our decoder on the first half and tested on the remaining half. Independent runs of the analysis were also run on arena maps divided into 16, 25, 36, or 49 spatial bins. Here, we again found evidence for increased decoding performance in Control MEC neurons versus APP KI MEC neurons (25 spatial bins) (**Supplemental Figure 8B, F**). For each session, position and speed decoding error ratios were consistently increased in Control MEC neurons versus APP KI MEC neurons, indicating a larger improvement from chance (Position: S1, *p*<0.001; S2, *p*<0.001; S3, *p*<0.05; S4, *p*<0.001) (Speed: S1, *p*<0.001; S2, *p*<0.01; S3, *p*<0.01; S4, *p*<0.001). This was also the case when we divided the arena space into 36 spatial bins (**Supplemental Figure 8C, G**), suggesting that the drop in performance is possibly driven by an overall reduced encoding reliability in APP KI subjects. For conditions in which the arena space was divided into 16 or 49 spatial bins, significant and non-significant effects were found, with overall trends consistent with increased decoding performance in Control MEC neurons vs APP KI MEC neurons (**Supplemental Figure 8A, D, E, H**).

### Aged APP KI mice exhibit neuronal hyperactivity in the MEC

Poor spatial coding during exploration of novel and familiar environments in MEC of APP KI mice may be linked to Aβ-associated dysregulation in neuronal firing activity. Indeed, neuronal hyperactivity and network dysfunction have recently been reported in this APP KI mouse model (Nakazono et al., 2017; Johnson et al., 2020; Takamura et al., 2021; Funane et al., 2022; Brady et al., 2023; Inayat et al., 2023; Yao et al., 2023), building on previous work demonstrating network hyperactivity in transgenic mouse models of Aβ pathology (Palop et al., 2007; Busche et al., 2008; Busche et al., 2012; Sanchez et al., 2012; Rodriguez et al., 2020).

To examine whether 18-month APP KI mice exhibit neuronal hyperactivity at the single unit level, we calculated the firing rates of individual APP KI neurons sampled from multiple Session 1 recordings across different recording blocks and compared them to sampled neurons from age-matched Control mice (for a detailed breakdown, please see Materials & Methods and **Supplemental Data 1**). A total of 1,142 MEC neurons were analyzed from 15 mice (Control, n=533 single units, n=7 mice; APP KI, n=609 single units, n=8 mice), with a minimum of 51 neurons sampled per mouse. Frequency histograms of the speed-filtered (3-100 cm/sec) AVG FRs revealed a larger proportion of hyperactive neurons (>5Hz: 14.12%) in APP KI mice versus Control mice (>5Hz: 6.75%) (**Figure 5A**). The proportion (%) of neurons exhibiting hyperactive (>5Hz) and normal (<5Hz) firing rates was then calculated per animal as its contribution towards the group’s total sample population. We found that APP KI mice contributed a greater proportion of hyperactive MEC neurons to their sampled population compared to Control mice (*p*<0.05), with no differences in the contribution of normal firing neurons between groups (**Figure 5D-E**). We also compared the cumulative frequency distributions of the collective firing rates for APP KI and Control mice. A two-sample Kolmogorov-Smirnov test revealed a shift towards increased firing rates in neurons from APP KI mice versus Control mice (*p*<0.001) (**Figure 5B**). This hyperactive phenotype was also evident upon calculating average firing rates per mouse from individual MEC neurons and then comparing the group mean values. Firing rates of MEC neurons in APP KI mice were significantly increased compared to Control mice (*p*<0.01) (**Figure 5C**).

**Figure 5.**
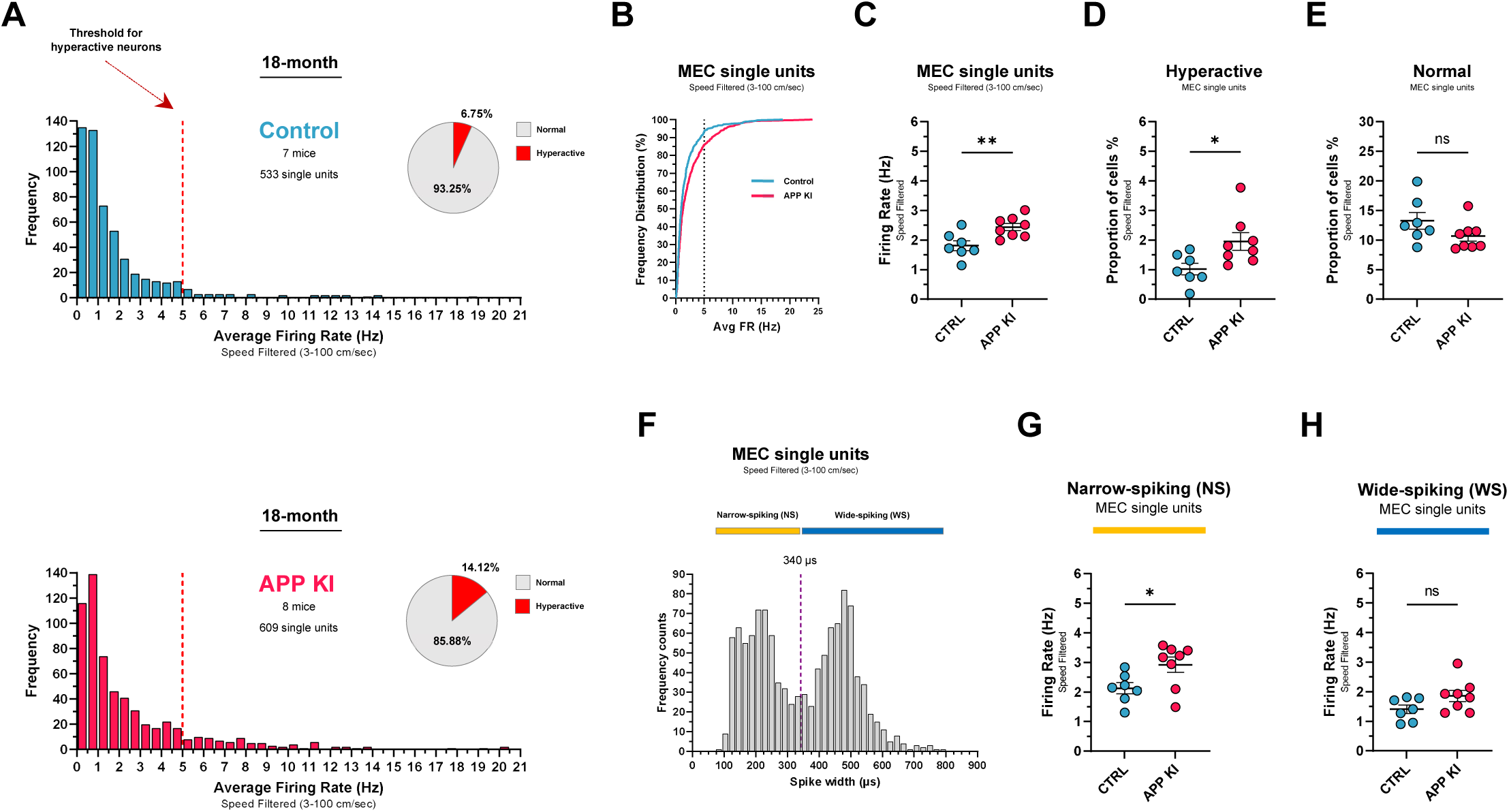
Aged *APP ^NL-G-F/NL-G-F^*mice exhibit subtle single unit hyperactivity in the medial entorhinal cortex. *In vivo* multielectrode recordings were performed in the medial entorhinal cortex (MEC) of 18-month APP KI mice and non-transgenic, age-matched C57BL/6J control mice as they freely explored open field arenas. A total of 1,142 MEC single units were collected from these mice (APP KI: Total, n=609 single units; n=7 mice; Control: Total, n=533 single units; n=8 mice). ***A***. Frequency distributions of the average firing rates for Control mice (top panel) and APP KI (bottom panel) after applying a min-max speed filter (3-100 cm/sec). Bins, 0.5 Hz. The dashed vertical line at 5 Hz indicates the threshold used to identify hyperactive single units. 14.12% of APP KI MEC single units were above threshold compared to 6.75% in Control. ***B*.** Cumulative frequency distributions of the collective single unit firing rates for both groups. Single units from APP KI mice exhibit a shift in distribution towards higher firing rates versus Control. Two-sample Kolmogorov– Smirnov test: APP KI versus Control: D = 0.1265, p < 0.001. The dashed vertical line indicates threshold for hyperactivity. ***C*.** Mean firing rates for APP KI mice were greater than Control. Unpaired t-test: *t* (13) = 3.099, *P* = 0.0085. ***D-E*.** The proportion of hyperactive (>5.0 Hz) MEC single units is greater in APP KI mice than Control. Unpaired t-test: *t* (13) = 2.527, *P* = 0.0252. No group differences were detected in the proportion of normal (<5.0 Hz) single units. Unpaired t-test: *t* (13) = 1.644, *P* = 0.1242. ***F*.** All speed-filtered MEC single unit firing rates were binned and plotted as a function of waveform spike width. The bimodal distribution in the frequency histogram was split at ∼340 μm spike width (dashed line) to delineate putative interneurons (NS, narrow-spiking units; yellow) from putative excitatory neurons (WS, wide-spiking units; blue). ***G-H*.** Mean firing rates of NS single units were greater in APP KI mice than Control. No group differences were detected in firing rates of WS single units. NS analysis - Mann-Whitney U test: *U* = 10, z = −2.025 *P* = 0.0401. WS analysis - Unpaired t-test: *t* (13) = 1.836, *P* = 0.0893.

To examine cell-type-specific firing patterns in our single unit dataset, we plotted cell frequency as a function of the neuronal waveform’s averaged spike width (μs) (**Figure 5F**) (Quirk et al., 2009; Rodriguez et al., 2020). The resulting bimodal distribution of the binned cell population allowed us to separate Narrow-Spiking (NS) cells from Wide-Spiking (WS) cells. Previous studies indicate that NS cells and WS cells likely correspond to putative interneurons and putative excitatory cells, respectively (Bartho et al., 2004; Quirk et al., 2009). A total of 575 NS cells and 567 WS cells were identified from our MEC recordings (NS, 50.35% versus WS, 49.65% of total neurons). APP KI mice exhibited increased neuronal firing rates in NS cells compared to Control mice (*p*<0.05) (**Figure 5G**). However, no differences were detected between groups in WS cells (**Figure 5H**).

Speed-filtered MEC oscillatory activity was also analyzed and compared across genotype in this dataset (**Supplemental Figure 9**). Similar to cross-session LFP analysis (**Supplemental Figure 3**), we did not detect group differences in percentage power of filtered LFP signals in the delta (1-3 Hz; *p*>0.05), theta (4-12 Hz; *p*>0.05), beta (13-20 Hz; *p*>0.05), low gamma (35-55 Hz; *p*>0.05), or high gamma (65-120 Hz; *p*>0.05) frequency ranges between APP KI and age-matched Controls.

Together, these results suggest that advanced Aβ pathology is associated with a mild hyperactive phenotype in MEC of aged APP KI mice that appears to be driven by putative interneurons.

## Discussion

The progressive loss of cognitive function in AD is accompanied by a pervasive increase in behavioral and psychological disturbances, including (but not limited to) agitation, restlessness, pacing, and wandering (Teri et al., 1988; Ballard et al., 2009). Spatial disorientation accompanied by wandering behavior has been reported in up to 65% of AD patients residing in assisted-living facilities, and is a chief concern to patient care providers and family members deciding on institutionalization (Kasper and Shore, 1994; Logsdon et al., 1998). While an increased tendency to wander and become lost is associated with mid- to late-stage AD, several studies have reported evidence for navigation and spatiotemporal orientation deficits early in disease progression (aMCI) and in individuals at genetic risk for AD (*APOE-*ε*4* carriers) (Parizkova et al., 2018; Coughlan et al., 2019; Bierbrauer et al., 2020; Lim et al., 2023; Newton et al., 2024; Peters-Founshtein et al., 2024). These reports suggest that path integration deficits may serve as strong indicators of incipient AD and warrant increased preclinical investigation into the neural mechanisms underlying pathology-associated degradation of spatial coding in the brain. For excellent reviews on the topic, please see (Coughlan et al., 2018; Segen et al., 2022; Igarashi, 2023).

In recent years, significant advances in our understanding of AD pathophysiology and its potential links to cognitive dysfunction have been made, largely through utilization of conventional transgenic mouse models that rely on overexpression of specific transgenes to recapitulate features of AD clinicopathology. However, transgenic mouse models that rely on overexpression of FAD-linked genetic mutations do not faithfully mimic the temporal or cell-type specific expression patterns that occur under normal physiological conditions. Thus, a number of phenotypic differences can occur across transgenic mouse models of Aβ and/or tau pathology, including age of onset for pathology, regional deposition and degree of pathology, as well as extent and severity of cognitive deficits. For excellent reviews on this topic, please see (Jankowsky and Zheng, 2017; Watamura et al., 2022).

The advent of human amyloid precursor protein (*App*) knock-in mouse models, wherein humanized Aβ is targeted to the endogenous murine app locus, was intended to circumvent potential confounds in transgenic mouse lines by preserving physiologic expression of hAPP (Saito et al., 2014). Several lines (e.g. *APP ^NL^, APP ^NL-F^, APP ^NL-G-F^*) were originally described, each exhibiting unique pathological Aβ profiles and behavioral deficits dependent on the mutations carried (Saito et al., 2014; Masuda et al., 2016). For instance, the *APP ^NL-G-F^* knock-in model stabilizes Aβ oligomers and protofibrils, leading to early Aβ plaque deposition throughout the brain. Aβ plaques begin around 3-months and appear near saturated in hippocampus and cortex at ∼7-months. *APP ^NL-G-F^* mice with significant Aβ pathology (6-13 months) exhibit deficits in context discrimination and spatial learning and memory, as well as impaired remapping in CA1 place cells and MEC grid cells (Johnson et al., 2020; Jun et al., 2020). Motivated by these findings, we chose to investigate spatial information coding in the MEC of 18-month *APP ^NL-G-F^* (APP KI) mice during free exploration of familiar and novel contextual environments in a two-day recording paradigm (**Figure 1A-B**). This allowed us to assess spatial information quality in individual MEC neurons tracked over hours long time intervals and quantify the stability of spatial coding features in MEC neurons across environments, all within a mouse model exhibiting significant Aβ pathology.

We found that APP KI MEC neurons encoded relatively poor spatial information in each session across the recording block, regardless of whether the mice had previously been exposed to the contextual environment or not (**Figure 1D-F**). Compared to Control MEC neurons, cross-session correlations of SI scores from APP KI MEC neurons revealed information coding instability early in the block (**Supplemental Figure 2A-B**), with later cross-session correlations trending towards increased instability (**Supplemental Figure 2C-I**). These data suggest that MEC neurons in APP KI mice encode minimal spatial information about a novel environment that does not improve over repeated exposures to the context, and that low-quality spatial information coding is inconsistent across familiar environments. We then examined single unit ratemaps from a larger dataset of recording blocks per mouse and found that MEC grid-cell-like representations were essentially absent in aged APP KI mice (**Figure 2A-B**), replicating a previous finding in relatively younger APP KI animals (7-13 months) (Jun et al., 2020). We also recorded border cells and head-direction cells from our aged mice. While head-direction cells recorded from mice in both groups maintained strong and stable directional tuning across sessions (**Figure 2E-F**), border cells recorded from aged APP KI mice exhibited increased instability in border firing preference and spatial tuning across sessions (**Figure 2C-D**). This instability did not appear to be dependent on any key session transition in the recording block, which included the first re-exposure to a given context (S1 > S2), the Day 1 afternoon to Day 2 morning transition (S2 > S3), or on the exposure to a novel context in the block (S3 > S4). Representative APP KI MEC border cells exhibited spatial remapping at different session transitions that included rotating border firing preference and sometimes forming or losing spatial tuning entirely (**Figure 2D**).

Disturbed grid cell periodicity in pathological Aβ-generating mouse models may be linked to deficits in MEC speed coding and improper integration of self-motion cues, leading to an overreliance on associations with geometric boundaries during navigation (Ying et al., 2023). Indeed, encountering environmental boundaries has been shown to correct spatial errors in grid cell firing (*i.e.* cohesive drift of firing fields) that accumulate relative to time and distance traveled since the last detected boundary (Hardcastle et al., 2015). In our paradigm, aged APP KI mice spent less time exploring the arena boundaries and more time in the arena centers than control mice (**Supplemental Figure 5F**). This difference in exploratory behavior may have contributed to increased error accumulation in the grid code of APP KI animals, as there was less opportunity for drift-based error correction. But what happens when boundary information coding itself is compromised? Compared to control mice, border cells in APP KI mice showed increased instability in firing field locations and spatial tuning across recording sessions, regardless of whether the mouse had previously been exposed to the environmental context or not (**Figure 2D**). This instability may serve as an additional source of uncertainty in judging contextual familiarity, reducing the utility of using this allocentric navigation strategy.

Neuronal activity patterns and spatial maps generated in the EC and hippocampus can undergo experience-dependent alterations known as ‘remapping’ (Muller and Kubie, 1987; Fyhn et al., 2007; Colgin et al., 2008). This flexibility in remapping of activity patterns can involve a change in firing rate only (rate remapping) or a change in firing rate accompanied by shifts in firing fields (global remapping) (Bostock et al., 1991). Remapping allows for dynamic encoding of various spatial representations under different environmental conditions and is thought to support episodic memory. However, stability in firing rates and/or firing fields within unchanged environments can also support episodic memory by generating consistent and reliable spatial representations, which could confer a degree of predictability across spatial maps. This may in turn strengthen the memory of a revisited environment and its conditions, possibly making navigation more efficient.

In this study, the stability of MEC spatial representations (*i.e.* ratemap firing densities) in aged mice was assessed using the whole map EMD, a metric based on optimal transport theory that was developed to circumvent the limitations of traditional ratemap correlation methods used in remapping studies (Aoun et al., 2023). The whole map EMD quantifies the minimum cost required to transform the 2D firing rate distribution of one whole ratemap (the origin ratemap) into another (the target ratemap), and is especially useful when dealing with non-linear transformations that feature changes in the shape and dispersion of firing fields. We calculated whole map EMD values for all matched MEC units in our recording block dataset, concentrating on session comparisons where mice were either re-exposed to a previously visited contextual environment (Context A: S1 vs S2, S2 vs S3) or exposed to a novel contextual environment (Context B: S3 vs S4). Reference distributions were then made using mismatched MEC units for each animal and session comparison to transform the raw, unbounded EMD values into bounded (0,1) quantile values for fair comparison (example reference distributions are shown in **Figure 3B-C**). Finally, matched MEC units were classified as spatial cells if their true SI scores exceeded the 90^th^ percentile in a distribution of shuffled SI scores (**Figure 3E**). A mixed-effects beta regression model was then run with genotype and spatial cell identity (isSpatial) as independent variables.

As predictors of ratemap instability, genotype and spatial cell identity had strong multiplicative effects on EMD quantile values for each session comparison in our recording paradigm (**Figure 3F**). Moreover, we found that median quantile values were larger in spatially-tuned APP KI MEC units versus control units during the first and second re-exposure to Context A, but not during exposure to a novel Context B (**Figure 3G**). These data suggest that MEC spatial cells in APP KI mice were several times more likely than controls to exhibit unstable spatial representations across the four-session recording paradigm, with the strongest effects occurring during exploration of familiar rather than novel contextual environments. Similar results were found when using an alternative shuffling method to identify MEC spatial cells in our mice (**Supplemental Figure 6**). Interestingly, previous work in mouse models of AD pathology has shown that repeated exposure to a once novel environment does little to improve stability in spatial representations (Zhao et al., 2014; Mably et al., 2017; Broussard et al., 2022). This would indicate that remapping events occur more often than expected in mice exhibiting Aβ and/or tau pathology, which likely contributes to the spatial learning and memory deficits observed in these models.

Across the population of MEC neurons recorded in our paradigm, spatial representations for position and speed were strongest in 18-month Control mice compared to age-matched APP KI mice (**Figure 4**). Control MEC neurons exhibited higher decoding performance for position and speed in both cross-session and within-session conditions, indicating stable spatial coding in and across familiar and novel contexts (**Supplemental Figure 7-8**). Conversely, spatial coding at the population level was disrupted in aged APP KI mice. APP KI MEC neurons showed poor decoding performance in cross-session conditions, suggesting a lack of coding generalization in a novel context (S4) as well as in familiar contexts (S2, S3). Interestingly, we also found that APP KI MEC neurons exhibited poor decoding performance in within-session conditions, likely reflecting a lack of stable spatial information coding throughout individual recording sessions (**Supplemental Figure 8**). Thus, it is not surprising that spatial coding features did not generalize across sessions in aged APP KI mice relative to Control mice.

The accuracy of our decoder was not particularly high for mice in either group, which we attributed to subject variability in exploratory behavior during free exploration of open field arenas. Overall behavioral performance metrics did not differ by genotype (**Supplemental Figure 5B-D**), though we did find that APP KI mice explored the arena centers more than Control mice in S1 and S3 (morning sessions) (**Supplemental Figure 5F**). Thus, while arena coverage and total distance traveled per session may not have been different between groups, the distribution of exploratory coverage certainly differed between subjects. This can be readily observed from the individual trajectory maps that make up the recording blocks (**Supplemental Figure 4**). To remedy this, we subsampled all sessions in the block so that they matched the minimum coverage among sessions for each spatial bin per condition. This ensured that all sessions had the same number of neural patterns per spatial bin, making it possible to fairly compare decoding performance between them. Bins were filtered for a minimum occupancy threshold of 100 samples and 100 samples from each bin were included to create balanced samples. Nonetheless, decoding accuracy in our study could perhaps have been improved with the use of a linear track, where behavioral variability can be more tightly regulated.

As previously discussed, *App*-KI second-generation mouse models were developed to circumvent confounding physiological consequences related to APP overexpression (Saito et al., 2014; Masuda et al., 2016; Watamura et al., 2022). One seemingly contentious phenotype observed in first-generation AD mouse models was that of increased nonconvulsive epileptiform-like activity in cortex and hippocampus, characterized predominantly by transient, large amplitude discharges reminiscent of epileptic interictal spikes (IISs) (Palop et al., 2007; Minkeviciene et al., 2009; Sanchez et al., 2012; Kam et al., 2016; Soula et al., 2023). The presence of IISs and other interictal epileptiform-like discharges (*e.g.* spike-and-wave, sharp wave, *etc.*) are thought to reflect disrupted excitatory/inhibitory balance in the brain that manifest as neuronal network hyperexcitability. Though these electrographic events in Aβ-generating mouse models bare striking similarity to events observed in studies of early-stage AD (Vossel et al., 2013; Vossel et al., 2016; Lam et al., 2017; Lam et al., 2020), preclinical reports have also suggested that APP overexpression during postnatal development is responsible for epileptiform-like abnormalities and not the overproduction of Aβ (Born et al., 2014; Kreis et al., 2021). However, second-generation *APP ^NL-G-F^* mice with no APP overexpression also exhibit interictal epileptiform-like discharges (8-12 months), in addition to neuronal hyperactivity and network dysfunction (3-16 months) (Nakazono et al., 2017; Johnson et al., 2020; Takamura et al., 2021; Funane et al., 2022; Brady et al., 2023; Inayat et al., 2023; Yao et al., 2023; Rajani et al., 2024). Recent *ex vivo* studies that record in *APP ^NL-G-F^* and *APP ^NL-F^* brain slices have implicated deficits in synaptic signaling and dysregulated homeostatic plasticity responses as potential drivers of aberrant activity in these models (Petrache et al., 2019; Calafate et al., 2023; Bonzanni et al., 2024). Together, these findings indicate that Aβ-associated disturbances in individual neurons subsequently lead to broader circuit and network dysregulation.

*In vivo* multielectrode recordings were used in our study to sample the activity of hundreds of MEC neurons from 18-month APP KI mice and age-matched controls, revealing a mild hyperactive phenotype driven by a subset of NS cells (**Figure 5**). To this end, we first generated a dataset containing >50 MEC neurons per mouse by combining units from multiple Session 1 recordings across several recording blocks (**Supplemental Data 1**). This allowed us to analyze MEC neuronal activity at the genotype (MEC units/group) and subject level (MEC units/mouse). Importantly, all Session 1 recordings used to analyze MEC activity per mouse were sampled from MEC positions ≥ 50μm from each other to minimize sampling any unit twice. Examining the frequency counts of binned AVG FRs for all units in each group revealed a small population of units that diverged at 5 Hz, with the APP KI group exhibiting an approximately two-fold increase in the number of ≥5 Hz units versus the control group (APP KI, 14.12%; Control, 6.75%) (**Figure 5A**). Comparing the cumulative frequency distributions of neuronal AVG FRs per group revealed a shift towards increased firing rates in APP KI MEC neurons versus control (**Figure 5B**). These findings were supported by further analysis at the subject level, with APP KI mice exhibiting increased firing rates versus control and contributing a higher proportion of hyperactive neurons to the recorded MEC population (**Figure 5C-E**). Finally, segregating MEC units based on their spike widths revealed increased firing rates in putative inhibitory interneurons (NS units) in aged APP KI mice versus control, with no effect of genotype on putative excitatory cells (WS units) (**Figure 5F-H**).

Several factors may underlie the observed increase in firing rates of MEC NS units sampled from our aged APP KI mice. One possibility may be a decrease in inhibition of MEC interneurons via disruptions in GABAergic signaling originating in the medial septum. Gonzalez-Sulser et al (2014) have shown that the medial septum sends monosynaptic inhibitory projections primarily to interneurons in the MEC, and previous work in various mouse models of AD pathology has demonstrated functional and anatomical remodeling of medial septum-hippocampal GABAergic projections (Rubio et al., 2012; Gonzalez-Sulser et al., 2014; Soler et al., 2017; Wander et al., 2023). Thus, it may be possible that Aβ pathology-associated deficits in GABAergic signaling along septocortical projections disinhibit MEC interneurons, leading to a subtle but detectable increase in interneuron firing rates in aged APP KI mice. A second possibility considers the unique susceptibilities of inhibitory interneuron subtypes to AD pathology. For example, somatostatin-positive (SST) and parvalbumin-positive (PV) interneurons are reported to exhibit distinct vulnerabilities at different stages of AD progression, though this may be brain region and cell layer specific (Petrache et al., 2019; Hijazi et al., 2020; Sanchez-Mejias et al., 2020; Morrone et al., 2022; Hijazi et al., 2023; Almeida, 2024; Gabitto et al., 2024; Goettemoeller et al., 2024; Kampmann, 2024). SST interneurons appear to be selectively vulnerable during the early stages of AD, with strong evidence highlighting their loss or dysfunction as an early hallmark of disease progression (Beal et al., 1985; Almeida, 2024; Gabitto et al., 2024). In contrast, PV interneurons seem to remain relatively unaffected until the later stages of AD, once pathology has progressed significantly (Hof et al., 1991; Petrache et al., 2019; Gabitto et al., 2024). Although we cannot identify interneuron subtypes from extracellular recordings, the hyperactive phenotype observed in MEC NS units in APP KI mice might reflect the differential susceptibility of certain interneurons to advanced Aβ pathology. Finally, the hyperactivity in putative interneurons we observed in MEC of aged APP KI mice may represent a late-stage compensatory response to counteract hyperexcitability in neighboring principal neurons affected by Aβ accumulation. Aberrant excitatory activity likely rose and peaked prior to 18-months of age, leading to remodeling of inhibitory circuits in MEC and increased firing rates in local interneurons (Palop et al., 2007; Johnson et al., 2020). So, while one might have expected a greater degree of hyperactivity and network dysfunction in our aged APP KI mice given the extensive Aβ deposition throughout the brain, the observed hyperactivity in NS units was relatively mild, suggesting a potential decline from an earlier, more pronounced hyperactive state that involved excitatory neurons. We speculate that this potentially residual hyperactivity in putative interneurons, while mild, may still play a significant role in disrupting spatial coding in the APP KI mouse line. Our findings underscore the importance of further research into interneuron dysfunction in mouse models of AD pathology and their contribution to information coding deficits.

In conclusion, our experimental results in aged APP KI mice suggest that advanced Aβ pathology interferes with spatial information coding in MEC during active exploration of contextually novel and familiar environments. Aβ pathology was associated with relatively poor spatial information quality in individual MEC neurons and increased instability in spatial maps of the environment, regardless of contextual novelty or familiarity. Aβ pathology was also associated with deficits in position and speed coding at the neuronal population level in each session of the recording paradigm. Finally, a relatively higher proportion of hyperactive MEC neurons were recorded in aged APP KI mice versus control mice, and this appeared to be driven by putative inhibitory interneurons in the population sample. An important step forward in this research area will be the continued utilization of knock-in mouse lines that preserve the native expression pattern of genes carrying disease-associated mutations (Saito et al., 2014; Xia et al., 2022; Watamura et al., 2023; Morito et al., 2024). Knock-in lines offer several advantages over traditional mouse lines that rely on transgene overexpression, and are especially useful for studies that probe pathology-associated neuronal circuit dysfunction and cognitive impairment.

### Limitations of the study

It is important to acknowledge several limitations of the study design and methodology described here that may impact the interpretation of the results. First, the recording blocks chosen for group analysis were selected from several blocks recorded per mouse over several weeks. This may have led to some difference in experience with handling and recording procedures in some mice versus others. However, we did not find genotype differences in overall locomotor activity within the selected recording blocks (**Supplemental Figure 5B-D**), suggesting that individual mouse differences in experience with the recording procedures had little to no effect on behavioral measures in the four-session block. Importantly, the presence of spatial cells (*e.g.* grid, border, head-direction, *etc.*) was not a criterion for the selection of recording blocks taken from each mouse. Rather, we attempted to balance the site of tetrode recordings across mice when selecting blocks for analysis, taking into consideration the quality of LFPs and single unit isolation, as well as ability to match single units across sessions by averaged waveforms on four channels. Nonetheless, we acknowledge the difficulty in matching precise recording locations across individual mice, and so we report a range of tetrode depths per group. The range of MEC recording depths for the recording block dataset was 950μm to 1700μm below dura for Controls and 1050μm to 1500μm below dura for APP KI mice, which we confirmed *post hoc* via histology (**Supplemental Figure 10**).

Second, we acknowledge that remapping events might have been more effectively triggered by altering the shape of the arena in the fourth and final recording session of the block. When designing the experimental paradigm, we chose to use 50cm x 50cm square arenas throughout the four-session blocks to keep the arena dimensions uniform for remapping (EMD) and decoding analyses across sessions. Contextual novelty was achieved by assembling square arenas with unique wall patterns using panels with high-contrast visual cues (**Supplemental Figure 1**). In doing so, we ensured that new combinations of arena contexts were used for each series of recording blocks per mouse. However, we cannot assert that the mice did not experience some familiarity with what we considered to be ‘novel’ contexts, or that mice experienced familiarity at all following repeated exposures to Context A in each block (S1 thru S3). In addition, it is also possible that mice may have attended to cues other than the arena walls during recording sessions (*e.g.* source of dim lighting in the recording room, sound localization of white noise machine, odor localization, *etc.*), which could have provided stable spatial references and influenced remapping events.

Finally, we acknowledge that our findings in APP KI mice are limited by the inclusion of a single age group (18-month), and that the addition of younger age groups would be useful in establishing the onset and progression of spatial coding deficits in this mouse line. Initial project planning of this study included the investigation of MEC spatial coding in two age groups of APP KI mice: a young (3-4 months) group and an aged (16-18 months) group. We chose to examine and report on this advanced age group for two main reasons. One, this project was launched in August 2020 in compliance with Columbia University’s phased ramp-up of research activities following a ramp-down resulting from the COVID-19 pandemic in New York City. Initial experimental recordings were performed in a small sample of younger (3-4 months) APP KI mice and C57BL/6J control mice. However, a series of pandemic-related disruptions in the lab’s mouse breeding program impacted the APP KI line and prevented us from increasing sample sizes to make up a younger cohort. Additional pandemic-related research impediments experienced that year compelled us to focus research efforts on our aged cohort. Second, our results in 18-month mice build on previous findings demonstrating early Aβ-associated impairment of spatial cell remapping, entorhinal-hippocampal circuit dysfunction, and spatial learning and memory deficits in younger APP KI mice (Johnson et al., 2020; Jun et al., 2020; Funane et al., 2022). Thus, we would expect that EMD and spatial decoding analyses would also yield deficits in younger APP KI cohorts.

## Materials and Methods

### APP ^NL-G-F/NL-G-F^ mouse colony, experimental animals and ethics statement

The homozygous *App ^NL-G-F/NL-G-F^* knock-in (APP KI) mice used in these studies feature a humanized murine Aβ sequence containing three familial AD mutations: the Swedish (KM670/671NL), Beyreuther/Iberian (I716F), and Arctic (E693G) mutations, and have been described previously (Saito et al., 2014). Original homozygous *App ^NL-G-F/NL-G-F^* males and females were obtained via material transfer agreement (MTA) with Drs. Takashi Saito and Takaomi C. Saido (RIKEN Center for Brain Science, Japan). Homozygous mice were backcrossed to C57BL/6J mice (JAX Strain #: 000664) every 10 generations to reduce genetic drift. F10 or greater heterozygous *App ^NL-G-F/wt^* breeding pairs were then crossed to regenerate homozygous *App ^NL-G-F/NL-G-F^*mice, and the colony was refreshed in this way every 10 generations. All homozygous APP KI mice used in these studies were sampled randomly from available aged cohorts in our colony and genotyped prior to the experiments described. Age-matched C57BL/6J mice were used as controls for these studies. A total of 18 mice, including males and females, were used. Total mouse numbers per genotype were as follows: *App ^NL-G-F/NL-G-F^* mice (Total, n=10; Female, n=6; Male, n=4) and age-matched, non-transgenic C57BL/6J mice (Total, n=8; Female, n=5; Male, n=3).

All mice were housed in a temperature and humidity-controlled vivarium at Columbia University Irving Medical Center and maintained on a standard 12 hr light/dark cycle with food and water provided ad libitum. All animal experiments were performed during the light phase in accordance with national guidelines (National Institutes of Health) and approved by the Institutional Animal Care and Use Committee of Columbia University (IACUC Approval Number: AABF3553. The Animal Welfare Assurance number: A3007-01).

### Microdrive construction and tetrode implantation

Microdrives were constructed as described previously (Fu et al., 2017; Rodriguez et al., 2020). Briefly, custom-made reusable 16-channel microdrives (Axona, UK) were outfitted with tetrodes consisting of twisted, 25-μm-thick platinum-iridium wires (California Fine Wire, USA). Prior to surgery, the tetrodes were cut to an appropriate length and electroplated with Platinum Black plating solution (Neuralynx, USA) until individual channel impedances dropped within a range of 150-200 Kohms. Final impedances were recorded in sterile 0.9% saline solution.

For electrode implantation, mice were first weighed and then anesthetized with isoflurane (3%–4% for induction; 0.5%–3% for maintenance) using a multichannel VetFlo Traditional Anesthesia vaporizer (Kent Scientific, USA) and fixed within a stereotaxic frame (Kopf Instruments, Germany). An intradermal injection of bupivacaine (Marcaine, 2mg/kg) was then administered at the clipped surgical site, followed by an incision to expose the skull 5 min later. Jeweler’s screws were inserted into the skull to support the microdrive implant. A 2-mm hole was made on the right side of the skull at position 3.0–3.1 mm lateral to lambda and approximately 0.2 mm in front of the transverse sinus (right hemisphere). An additional screw connected with wire was inserted into the skull (left hemisphere), serving as a ground/reference. The prepared microdrive was then tilted at 6°–7° in the sagittal plane on a stereotaxic arm and the tetrodes lowered into the brain 1.0 mm from the surface (below dura). The microdrive ground wire was then soldered to the skull screw wire and the microdrive was secured with dental cement. Mice were then allowed to recover in a cleaned cage atop a warm heating pad until normal breathing and ambulation had returned before being transported to non-barrier housing for post-operative monitoring. Mice received Carprofen (5 mg/kg) prior to surgery and postoperatively to reduce pain, in addition to a subcutaneous sterile saline injection to aid in hydration. All implanted animals were housed individually to prevent damage to their microdrives and recording experiments began approximately 1 week from the time of surgery.

### Experimental design: recording blocks

*In vivo* recording sequences consisted of recording blocks made up of four individual recording sessions spread over two consecutive days, with two recording sessions performed each day (**Figure 1B**). Sessions 1 and 2 were recorded on Day 1 in the morning and afternoon, respectively. Sessions 3 and 4 were recorded in the morning and afternoon of the following day (Day 2). The inter-session interval per day was ∼4-6 hr and the time interval between days (Session 2 to 3) was ∼16-18 hr. Once a recording block was completed for a particular recording depth, tetrodes were advanced 50 to 100µm into the MEC to sample neuronal activity from a new population of neurons. All mice were allowed 48 hr before the next recoding block was performed.

As illustrated in **Figure 1B**, Session 1 is the first exposure to Context A (novel), in this case depicted as a square arena with black walls and a white cue card affixed to the North wall. Sessions 2 and 3 constitute re-exposures to Context A (familiar), while Session 4 is an exposure to a novel panel arrangement (Context B), depicted in **Figure 1B** as a square arena with white walls and a black cue card affixed to the East wall. The individual panels used to construct the walls (boundaries) of square open field arenas were white and black 3/16” sturdy foam board (50cm x 80cm) cut from larger one-piece boards (Staples, USA). Panels featured high contrast visual cues, though solid white or black panels with no cues were also used. 80cm white composite corner trim (90° angle) was used at each panel end to assemble a square arena. The variety of panels allowed construction of unique arena contexts for repeated *in vivo* recording sessions while maintaining arena dimensions. Arena dimensions once assembled: 50cm x 50cm x 80cm. All panel arrangements used in the experiments described here are found in **Supplemental Figure 1**.

The subject order for daily recordings was randomly assigned so that the same sequence of animals was never repeated across days, and every effort was made to balance the subject order by genotype and sex for each recording block. To begin a recording block (Day 1), mice were transported in their home cages to an isolated recording room within the housing facility and allowed to acclimate for ∼30 min. Once a square arena was configured (Context A) and a subject order was established for the day, mice were removed from their home cages, connected to the recording system, and allowed to freely explore the open field for 20 min. Motivated foraging behavior and open field coverage were encouraged by scattering chocolate crumbs throughout the arena. Once Session 1 was completed, mice were gently lifted out of the arena and disconnected from the recording system before being placed back into their home cages. Session 2 recordings were performed under the same experimental conditions and followed the same subject order. The following day (Day 2), the subject order was again randomized prior to recording. Session 3 recordings were performed under the same conditions and in the same arena as Day 1 recordings. Once completed, a novel contextual environment (Context B) was created by assembling a new arena with a panel configuration that differed from Context A. Session 4 recordings were then performed in Context B under the same conditions as prior recordings. The arena space was thoroughly cleaned between each recording session to minimize the influence of odors from a previously run animal. All recordings were performed under low light conditions (ambient room, ∼25 lux; center of arena, ∼8 lux) with background white noise (50db).

### In vivo recording and single unit analysis

Neuronal signals from our mice were recorded using the Axona DacqUSB system and described previously (Fu et al., 2017). Briefly, recording signals were amplified 10,000 to 30,000 times and bandpass filtered between 0.8 and 6.7 kHz. Spike sorting was performed offline using TINT cluster-cutting software and KlustaKwik automated clustering tool. The resulting clusters were then refined manually and were validated using autocorrelation and cross-correlation functions as additional separation tools. Single units with no undershoot in their waveform were discarded.

A total of 1,142 MEC neurons were collected across 15 mice for data shown in **Figure 5** (Control, n=533 single units (Total, n=7 mice; Female, n=5 mice; Male, n=2 mice); APP KI, n=609 single units (Total, n=8 mice; Female, n=4 mice; Male, n=4 mice)). This dataset was generated by sampling MEC neuronal activity from multiple Session 1 recordings across different recording blocks for each mouse. This approach allowed us to capture new populations of MEC neurons in each mouse while ensuring consistency in session sampling across animals. First, a minimum–maximum speed filter (3-100 cm/sec) was applied to remove artifacts due to LED jumps and single unit spikes that occurred during bouts of behavioral immobility. Next, quantitative measurements of cluster quality were performed, yielding isolation distance values in Mahalanobis space (Schmitzer-Torbert et al., 2005). Median isolation distance values were then calculated per mouse followed by comparison across genotype. No significant difference in cluster quality was detected between groups (median isolation distances: Control, 8.390; APP KI, 7.887: Mann-Whitney *U* test: *U* = 20, *P* = 0.3969). Average firing rates were calculated by dividing the total number of speed-filtered spikes by the duration of recording session (1,200 sec). A conservative threshold for single unit hyperactivity (5.00 Hz) was determined based on a previous finding in 16-month EC-Tau/hAPP mice (4.74 Hz) and hAPP mice (5.60 Hz) after examining the relative change in proportions of neurons in APP KI mice versus Controls (Busche et al., 2008; Rodriguez et al., 2020). Putative excitatory neurons (WS) were distinguished from putative interneurons (NS) in this dataset by first examining the frequency distribution histogram of pooled MEC single unit spike widths and then bisecting the binned waveform spike widths after the first modal distribution (Bartho et al., 2004; Quirk et al., 2009; Rodriguez et al., 2020). This resulted in a delineation cutoff at 340μs (**Figure 5F)**. A total of 575 NS cells and 567 WS cells were identified from our MEC recordings (NS, 50.35% versus WS, 49.65% of total neurons). For a detailed breakdown of the electrophysiology dataset, please see **Supplemental Data 1**.

A total of 226 MEC neurons across 15 mice were identified and tracked across sessions in recording blocks described in **Figures 1 thru 4, Supplemental Figures 1 thru 8, Table 1**, and **Supplemental Table 1**: Control, n=115 matched single units (Total, n=8 mice; Female, n=5 mice; Male, n=3 mice); APP KI, n=111 matched single units (Total, n=7 mice; Female, n=4 mice; Male, n=3 mice). ‘Matched’ neurons were any neurons that appeared in each session of the recording block and that could be matched based on clusters in 2D feature space and average waveform. ‘Non-matched’ neurons were any neurons that did not appear in all four sessions of the recording block or those that could not be reliably matched. A total of 126 MEC neurons were classified as Non-matched (Control, n=68 non-matched single units; APP KI, n=58 non-matched single units) and were not included in recording block analysis of tracked neurons.

### Animal performance

Behavioral analysis of animal performance in the open field arenas has been described previously (Rodriguez et al., 2020). Animal performance during in vivo recording sessions was assessed by tracking the position of an infrared LED on the head stage (position sampling frequency, 50 Hz) by means of an overhead video camera. The position data were first centered around the point (0, 0), and then speed-filtered where noted, with only speeds of 3 cm/second or more included in the analysis. Tracking artifacts were removed by deleting samples greater than 100 cm/second and missing positions were interpolated with total durations less than 1 second, and then smoothing the path using a moving average filter with a window size of 300 milliseconds (15 samples on each side). The total distance traveled in the arena (m), % of arena coverage, and average speed (cm/second) during exploration served as dependent measures of interest. For % of arena coverage, the processed position data was plotted and converted to black and white. Using the “bwarea” function in MATLAB, we calculated the total area traveled by the mouse using the binary image and then divided this area by the total area of the arena.

To analyze the amount of time mice spent exploring the center of the maze versus the boundaries, a 4×4 grid was imposed on the arena area and the resulting four centralized boxes were defined as the ‘Center’ zone (625cm^2^). The remaining twelve boxes corresponding to the arena borders were defined as the ‘Boundaries’ zone (1,875cm^2^). Center and Boundaries Zone occupancy (%) was then calculated by summing the time spent in boxes corresponding to each zone as a percentage of the total session duration (20 min). Trajectory maps from each session in the recording blocks are shown for each mouse along with the total distance traveled (m) and % arena coverage (**Supplemental Figure 4**). Mouse paths appear yellow on a dark background. A 4 x 4 grid (white lines) is overlaid onto the trajectory maps. Arena dimensions: 50cm x 50cm.

### Spatial information and spatial cell analysis

Spatial information content is a measure used to predict the location of an animal from the firing of a neuron. For each neuron, information content was calculated using Skaggs’ formula, which measures the amount of information carried by a single spike about the location of the animal and is expressed as bits per spike (Skaggs et al., 1993).

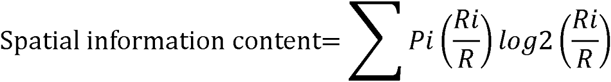

where i is the bin/pixel number, Pi is the probability for occupancy of bin i, Ri is the mean firing rate for bin/pixel i and R is the overall firing mean rate.

Firing ratemaps were computed by first making 64×64 unit 2D spatial histograms for occupancy and spike data. Both histograms were then smoothed using a 16×16 unit Gaussian Kernel with a standard deviation of 3.2 units. The smoothed spike map was then divided unit-wise by the smoothed occupancy map to obtain the final rate map. The Peak Firing Rate was defined as the rate in the bin with the highest rate in the firing rate map.

Grid, border, and HD cells shown in **Figure 2** were identified according to spatial firing characteristics described in published reports (Hafting et al., 2005; Langston et al., 2010; Bonnevie et al., 2013; Kropff et al., 2015; Fu et al., 2017). Briefly, units were classified as grid cells if they exhibited hexagonal firing patterns with well-formed firing fields in any recording block session. Border cells were identified by a characteristic firing field in the proximity of an environmental boundary, and HD cells were identified if units preferentially displayed tuning toward a particular direction in polar plots. Spatial cells shown as examples are from 5 Control mice and 7 APP KI mice. Grid, border, and HD spatial tuning properties were analyzed in Python using the open-source package *opexebo* provided by the Kavli Institute for Neuroscience in Trondheim, Norway (https://pypi.org/project/opexebo/) (Obenhaus et al., 2022). Grid, border, and HD scores were generated and appear at the top of ratemaps and polar plots in **Figure 2**. However, statistical analysis was not performed on spatial cells in this study due to an insufficient number of spatial cells collected.

For EMD analysis (**Figure 3**), SI scores were used to identify ‘spatial’ and ‘non-spatial’ MEC single units using two independent shuffling procedures: circular shuffling and firing field shuffling. For each procedure, spatial cells were defined as those units whose Session 1 SI scores exceeded the 90^th^ percentile of the shuffled SI score distributions created from Session 1 Control data (Total, n=115 units; n=8 mice). For circular shuffling, spike times and positions were misaligned by circularly shifting spike timestamps by a constant between 20 seconds and T-20 seconds, where T represents the duration of the recording sessions. This constant was randomly selected for each cell on each of 1,000 shuffles. Ratemaps were recomputed after circular shuffles and a distribution of shuffled information scores was created. A total of 114,038 circularly shuffled SI scores were used to create the distribution in **Figure 3E** from an original total of 115,000 computed scores. Notably, 962 circularly shuffled SI scores were discarded because they exceeded 2.0 SI (range: 2.0-3.99). 640 shuffled scores from mouse 1-28 (T1C1) were discarded, along with 322 shuffled scores from control mouse 1a23 (T2C10). For field shuffling, the positions of segmented firing fields were randomly shuffled within their respective ratemaps using a method described here (Grieves et al., 2021). This shuffling method essentially disrupts the original spatial order of firing fields while preserving as best as possible the internal structure of the fields and firing activity. 1,000 field shuffles were performed per unit and a distribution of field shuffled SI scores was created, yielding a total of 115,000 field shuffled SI scores. The 90^th^ percentile cutoff for the circularly shuffled SI distribution was 0.350168 and resulted in 63 spatial units total (Control, n=38 spatial units; APP KI, n=25 spatial units). The 90^th^ percentile cutoff for the firing field shuffled SI distribution was 0.3589145 and resulted in 61 spatial units total (Control, n=38 spatial units; APP KI, n=23 spatial units).

### Spatial remapping using the Earth Mover’s Distance

The Earth Mover’s Distance (EMD) was used to quantify the spatial dissimilarity between two rate map distributions in **Figure 3** (Aoun et al., 2023). The EMD can be thought of as the minimum cost required to transform one distribution into another, where the cost is proportional to the amount of “earth” moved and the distance it is moved. Here, “earth” represents the firing density and the distance represents the change in that density. To efficiently compute this distance on two-dimensional maps, the EMD was approximated using the sliced EMD (Bonneel et al., 2015). The sliced EMD is computed by projecting the rate map into many random one-dimension slices, and then computing the EMD on each of these slices using the one-dimensional closed-form solution:

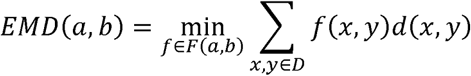

The average of the distances across multiple slices (10^3^) converges close to the true two-dimensional EMD.

We then computed an EMD metric (Wasserstein distance) across normalized firing rate distributions (ratemaps). The ‘whole map’ EMD distances bins spike counts across the full arena dimensions and is computed for each cell’s pair of rate maps on a given session to session comparison. To standardize the EMD remapping scores, we converted them into quantiles with animal and time interval specific counterfactual distributions. We defined each counterfactual distribution to be the EMD values from mismatched neurons across tetrodes for each animal in specific session to session comparisons (*i.e.* S1 vs S2, S2 vs S3, S3 vs S4). For each animal, and for a given session to session comparison, we computed the sliced EMD between every possible pair of incorrectly matched neurons from those two sessions. We then compared the EMD values from the correctly matched neurons to the distribution of incorrectly matched neurons, yielding quantiles. These quantiles represent estimates of the probability (observed proportions) that each correctly matched neuron would have a lower EMD (*i.e.* a more stable spatial representation) than randomly mismatched neurons from the same animal and the same session pair. It is important to note that the quantiles for coherent spatial representations that have been remapped can look similar to quantiles for non-spatial cells. Therefore, the quantiles only serve as a standardized measure of representational stability and not as a measure of spatial information.

To compare standardized remapping scores, we used mixed-effects beta regression models for each session transition. In addition to distinguishing between genotype, we categorized MEC single units as spatial cells or non-spatial cells using a binary ‘isSpatial’ variable. To do so, we misaligned spike times and positions by circularly shifting spike timestamps by a constant between 20 seconds and T-20 seconds, where T represents the duration of the recording sessions. This constant was randomly selected for each cell on each of 1,000 shuffles. Rate maps were recomputed after circular shuffles and a distribution of shuffled information scores was created. Given this distribution, each spatial information score was associated with a p-value. For a given cell, it was defined as spatial if at least one tested session had a significant spatial information score. Thus, our beta regression included three fixed effects and one random effect. The three fixed effects were: (1) genotype (control served as the reference state), (2) spatial cell status (reference was non-spatial), and (3) the interaction between (1) and (2). We included the animal ID as a random effect to control for correlations from samples within the same animal. We reported our results as odds ratios derived from the sum of the genotype, spatial cell and interaction effects. Our results thus quantified the impact of APP KI genotype on the stability of spatial representations in spatial cells across recording sessions.

### Spatial decoding using a support vector machine (SVM)

We assessed the relative spatial decoding performance between groups by training an ensemble of SVMs on single unit activity from each group. We assessed relative decoding performance for two target variables: (1) position, and (2) speed. For position, we divided the animal arena into 16, 25, or 36 spatial bins. For speed, we divided the distributions into between 4 and 6 bins. Each target variable and number of bins corresponded to a separate ‘run’ of the pipeline.

During a given run, we divided the dataset into single units from the APP KI group and the Control group. For each group, we trained an ensemble of SVMs using a classification strategy called One Versus One (OVO), where one model is trained per bin and then they are all combined in order to yield the final prediction. The prediction error for each bin was computed in centimeters by taking the average of the Euclidean distances from each predicted bin to the center of the true bin for each target bin. Next, for each bin, the mean error was compared to the mean error from the shuffled predictions within the same training run. The ratio of the true model error to the shuffled error was computed for each bin. Finally, the error ratios for all the bins from the Control group were compared to the bins from the APP KI group using the Wilcoxon signed-rank test (**Figure 4C, E**).

### Brain tissue fixation and slice preparation

At the end of our experiments, all mice were deeply anesthetized with a cocktail of ketamine/xylazine before being transcardially perfused with ice-cold 100mM phosphate-buffered saline (PBS) pH 7.4., followed by 10% formalin (Fisher Scientific, USA). The last recording position for each microdrive-implanted mouse was noted, followed by careful removal of the microdrive. Mouse brains were then harvested and incubated in 10% formalin overnight, followed by incubation in 30% sucrose until the brains sank to the bottom of a 15mL conical tube (all at 4°C). Following cryoprotection, brain hemispheres from microdrive-implanted mice were separated and frozen in O.C.T. mounting medium. The right hemispheres (implanted) were sliced into sagittal brain sections (40µm) using a Leica CM3050 S cryostat and stored in cryoprotectant at −20°C following immunostaining procedures. Left brain hemispheres (non-implanted) were stored at −80°C.

### Histology: Immunofluorescence and Nissl staining

Immunostained sections in **Figure 1** were collected from a naïve, age-matched set of *App ^NL-G-F/NL-G-F^* and C57BL/6J mice sampled from our colony. Briefly, free-floating horizontal brain sections (40µm) were rinsed in PBS and then permeabilized with 0.3% Triton X-100 (PBS-T). Sections were then blocked in 10% normal goat serum (NGS) in PBS-T for 60 min at room temperature, followed by overnight incubation at 4^0^C in primary antibodies diluted in 5% NGS in PBS-T. Primary antibodies used were anti-amyloid beta (Aβ) D54D2 (rabbit monoclonal, 1:1,000) (Cell Signaling, USA) and anti-NeuN (guinea pig polyclonal, 1:5,000) (Synaptic Systems, Germany). The next day, sections were rinsed in PBS-T and incubated in secondary antibodies diluted in 5% NGS in PBS-T. Secondary antibodies used were goat anti-rabbit IgG AlexaFluor 488 (1:500) (Life Technologies) and goat anti-guinea pig IgG Alexa Fluor 594 (1:500). Sections were then incubated in a working solution of Hoechst 33342 dye (5µg/mL) (Thermo Scientific, USA) to stain cell nuclei for 3 min at room temperature. Subsequent washes in PBS were followed by mounting the tissue onto Superfrost Plus slides and coverslipping using SlowFade gold anti-fade reagent (Life Technologies, USA). All slides were stored in the dark at 4^0^C until imaging.

To verify tetrode placement and reconstruct recording positions in our experimental mice (**Supplemental Figure 10**), sagittal tissue sections from implanted hemispheres were mounted onto gelatin coated glass slides and stained in a filtered solution of 0.1% Cresyl violet dissolved in distilled water. Slides were then coverslipped using DPX mountant and stored at room temperature until imaging.

### Imaging

Fluorescence and brightfield microscopy were performed using an Olympus BX53 upright microscope equipped with an Olympus DP72 12.5 megapixel cooled digital camera. Image files were saved locally on a Dell computer running the Olympus cellSens software platform, then transferred to a Hussaini lab Cyberpower PC for offline image processing in Fiji (Image J) (Schindelin et al., 2012).

### Statistical Analysis

Statistical analyses were performed in GraphPad Prism (version 10.4.0) and R (version 4.4.1). All datasets were tested for normality using the Shapiro-Wilk test. Datasets in which values were not modeled by a normal distribution were subjected to nonparametric statistical analyses. In Figure 1C, an unpaired *t*-test was used to compare the group means of matched MEC single units (%) tracked in our recording block. Correlation matrices were then assembled to visualize relationships between matched MEC single unit’s SI scores, speed-filtered AVG FRs, and speed-filtered Peak FRs in Figure 1D-E. The resulting Pearson’s correlation coefficients (r) were color scaled using the ‘magma’ colormap in GraphPad Prism (1.0, yellow; −1.0, black). Unpaired *t*-tests were used to compare the SI group means for each session in Figure 1F. A mixed-effects beta regression model was used to determine variable effects on the whole map EMD in Figure 3D and 3F. Summary statistics are shown in Table 1. Genotype and isSpatial variables, and their interaction, were considered fixed effects while the Subject ID was included as a random effect to control for correlated samples. The mixed effects models were computed using the glmmTMB R package. Diagnostic tests for dispersion, zero-inflation and outliers were computed using the DHARMa package with non-significant results for all tests. Mann-Whitney *U* tests were used to compare the group median quantile values in each session comparison shown in Figure 3E. Wilcoxon signed-rank tests were used to assess genotype differences in decoding error ratios for position and speed in Figure 4C, E. A two-sample Kolmogorov-Smirnov test was used to compare the distributions of speed-filtered AVG FRs in Figure 5B. Unpaired *t*-tests were used to compare group means in Figure 5C-E and 5H.

Welch’s *t*-tests were used to compare group means of Pearson’s correlation coefficients (*r*) for each session comparison in Supplemental Figure 2. Repeated measures two-way ANOVAs were used to calculate the effects of two independent variables (genotype x session) on the percentage of power values for individual oscillatory frequency bands in Supplemental Figure 3. A Mann-Whitney *U* test was used to compare group means in Figure 5G. A repeated measures two-way ANOVA was used to calculate differences in performance metrics (Total Distance, Arena Coverage, Avg Speed) over time in Supplemental Figure 5B-D. Correlation matrices of animal performance metrics are shown in Supplemental Figure 5E. The resulting Pearson’s correlation coefficients (r) were color scaled (1.0, white; −0.5, black). A three-way ANOVA was used to calculate the effects of three independent variables (genotype x session x zone) on the percentage of time mice spent in the arena zones across sessions in the recording block in Supplemental Figure 5F. As previously described for Figure 3, a mixed-effects beta regression model was used to determine variable effects on the whole map EMD in Supplemental Figure 6B. Summary statistics for this analysis appear in Supplemental Table 1. Mann-Whitney *U* tests were used to compare the group median quantile values in each session comparison shown in Supplemental Figure 6C-D. Wilcoxon signed-rank tests were used to detect genotype differences in Supplemental Figures 7-8. Unpaired *t*-tests were used to compare means in Supplemental Figure 9. *Post hoc* analyses were performed using Tukey’s multiple comparisons test where noted.

## Data and code availability

The source data used to generate all graphs will be included as **Supplemental Data 1** with this manuscript after peer review and acceptance for journal publication. Source data will be organized by individual sheets labeled as figures in an Excel file and made freely available on the lab’s GitHub page. The code used for data analysis will also be made freely available on the lab’s Github page once the article has been peer reviewed and accepted for journal publication. It can also be requested from the corresponding author prior to journal publication.

## Author Contributions

Conceptualization, G.A.R. and S.A.H.; Methodology, G.A.R., E.F.R., C.O.S., A.A., L.P., T.T.; Software, C.O.S., A.A., L.P., S.V.V.; Formal analysis, G.A.R., E.F.R., C.O.S., A.A., L.P.; Investigation, G.A.R., E.F.R., T.T.; Resources, G.A.R., C.O.S., A.A., L.P., S.V.V., S.F., S.A.H.; Data curation, C.O.S., A.A., L.P.; Writing – original draft, G.A.R.; Writing – review & editing, G.A.R., E.F.R., C.O.S., A.A., S.F., S.A.H.; Visualization, G.A.R., C.O.S., A.A., L.P.; Supervision, S.F., S.A.H.; Project administration, G.A.R., S.A.H.; Funding acquisition, G.A.R., S. F., S.A.H.

## Acknowledgments

The authors wish to thank Drs. Takashi Saito and Takaomi C. Saido (RIKEN Center for Brain Science, Japan) for providing the *APP ^NL-G-F^* knock-in mice used to establish the Hussaini lab colony at Columbia University Irving Medical Center. We also wish to thank Helen Y. Figueroa for maintaining the aged *APP ^NL-G-F^* and C57BL/6J mouse colonies used for this study. We wish to thank Dr. Fabio Stefanini for his role in the early conceptualization of the decoding analysis, and we wish to thank Dr. Radha Raghuraman and Dr. Alexandra Petrache for helpful discussions and comments regarding the manuscript. This work was supported by research grants from the National Institute on Aging (R01AG050425; R01AG064066) to S.A.H., (K01AG068598) to G.A.R., (RF1AG080818) to S.F., and the BrightFocus Foundation (A2019382F) to G.A.R. The funders had no role in the study design, data collection and analysis, decision to publish, or preparation of this manuscript.

---

**Supplemental Figure 1.**
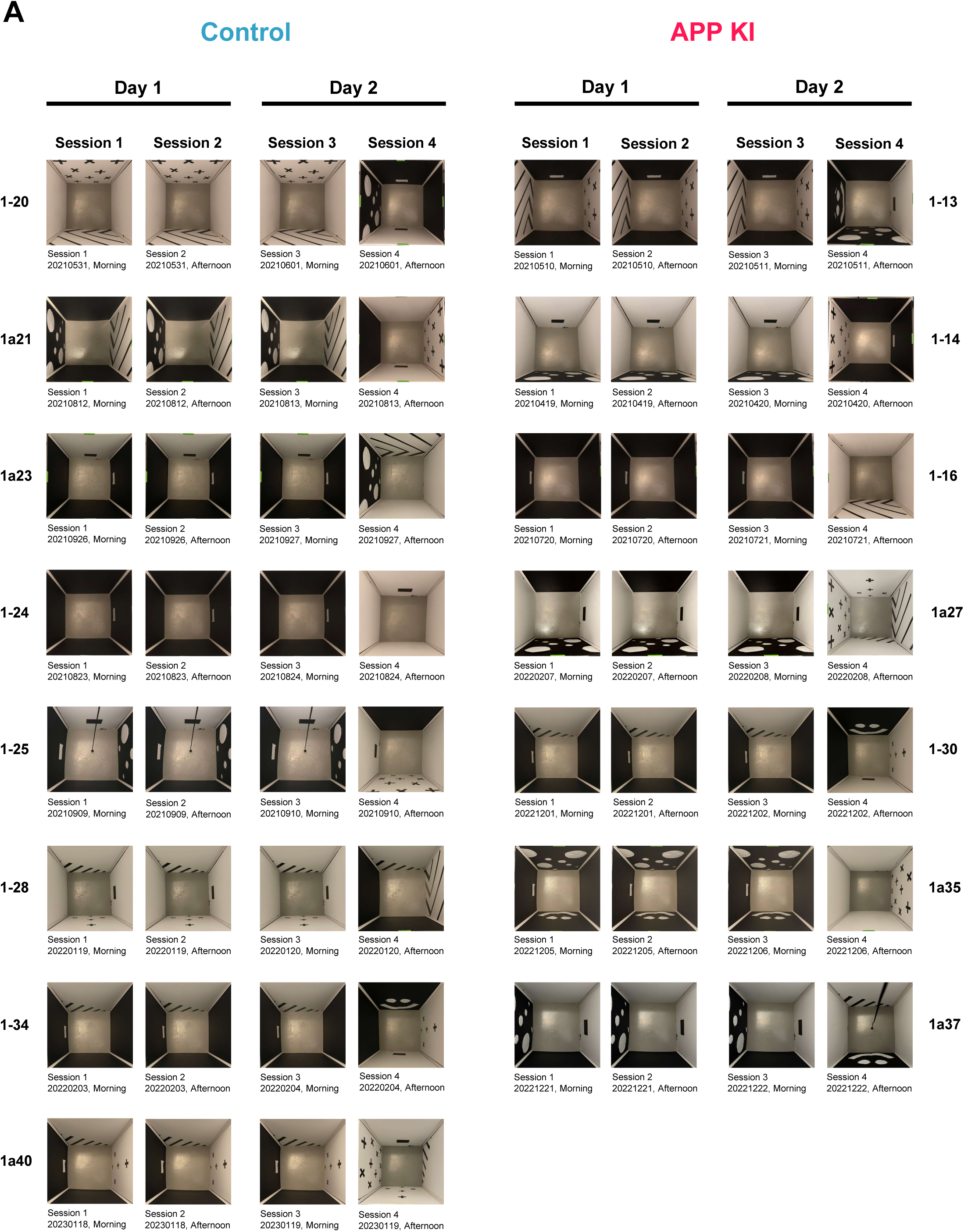
Recording block arena contexts. ***A*.** All panel arrangements that make up arena contexts used in experimental recording blocks are shown along with session date. Individual panels are white and black 3/16” sturdy foam board (50cm x 80cm) that feature various high contrast cues, though solid white or black panels were also used. 80cm white composite corner trim (90° angle) was used at each panel end to assemble a square arena. The variety of panels allowed construction of unique arena contexts for repeated *in vivo* recording sessions while maintaining arena dimensions. Arena dimensions once assembled: 50cm x 50cm x 80cm (LxWxH).

**Supplemental Figure 2.**
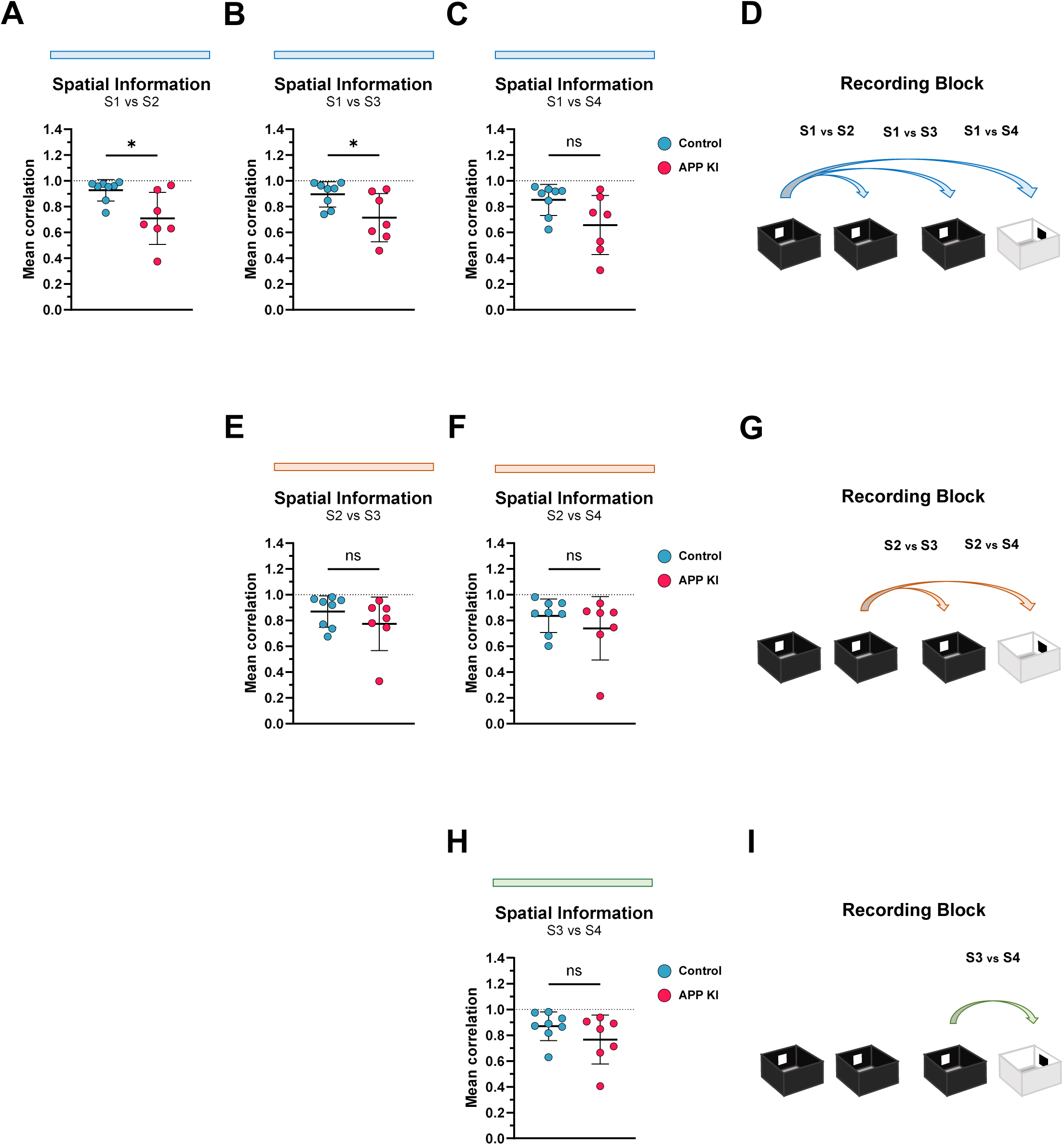
Recording block cross-session correlations of spatial information scores. Spatial information (SI) scores were calculated for each consistently tracked (‘Matched’) medial entorhinal cortex (MEC) single unit in our recording block. Pearson’s correlation coefficient (*r*) was then calculated for all single units per mouse based on session comparisons to assess informational stability across the recording block (Control, n=8 mice; APP KI, n=7 mice). For each session comparison, the group means were compared using Welch’s t-test. Overall, cross-session correlations of SI scores from Control MEC single units were quite strong, whereas corresponding correlations of SI scores from APP KI MEC single units were moderate. ***A-C*.** Genotype differences in mean Pearson’s *r* values were detected on the following session comparisons: Session 1 vs Session 2 (Control, 0.9267 ± 0.0833; APP KI, 0.7089 ± 0.2016) (*t*=2.667; *p*<0.05) and Session 1 vs Session 3 (Control, 0.8951 ± 0.0981.; APP KI, 0.7142 ± 0.1866) (*t*=2.302; *p*<0.05), but not Session 1 vs Session 4 (Control, 0.8521 ± 0.1204; APP KI, 0.6579 ± 0.2289) (*t*=2.013; *p*>0.05). ***D*.** An illustration of recording session comparisons for A-C. ***E-F*.** No genotype differences in mean Pearson’s *r* values were detected in Session 2 vs Session 3 (Control, 0.8684 ± 0.1215; APP KI, 0.7735 ± 0.2086) (*t*=1.057; *p*>0.05) or Session 2 vs Session 4 (Control, 0.8364 ± 0.1302; APP KI, 0.7393 ± 0.2456) (*t*=0.937; *p*>0.05) session comparisons. ***G*.** An illustration of recording session comparisons for E-F. ***H*.** No genotype differences in mean Pearson’s *r* values were detected in Session 3 vs Session 4 (Control, 0.8701 ± 0.1122; APP KI, 0.7664 ± 0.1893) (*t*=1.268; *p*>0.05). ***I*.** An illustration of the recording session comparison for H. Scatter plots and numerical values in figure legend represent the mean ± standard deviation of the Pearson’s *r* values per mouse in each group. * *p*<0.05.

**Supplemental Figure 3.**
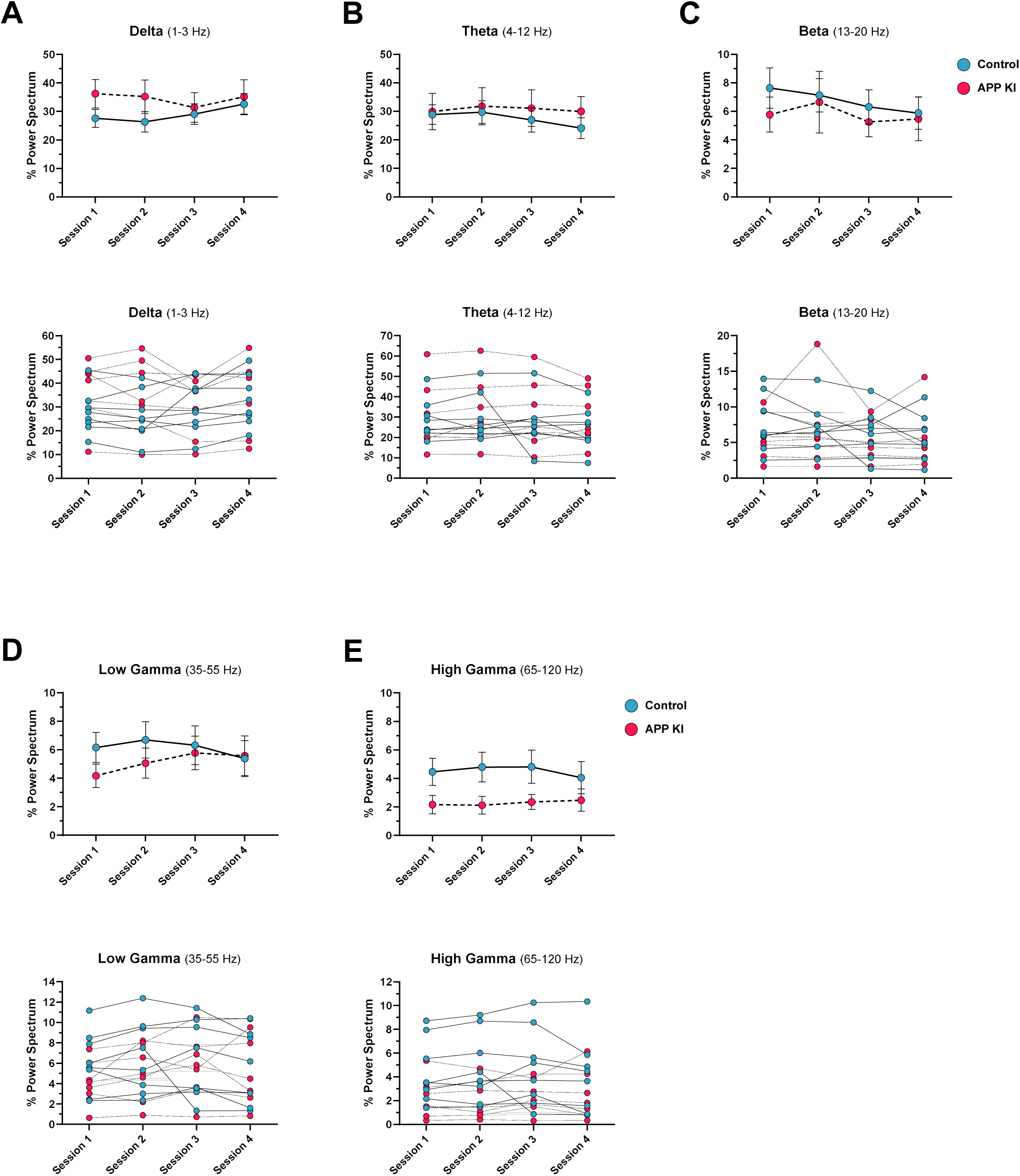
No effect of genotype on *in vivo* medial entorhinal cortex oscillatory activity throughout recording block sessions. Local field potentials (LFP) were recorded from the medial entorhinal cortex (MEC) of 18-month *APP ^NL-G-F/NL-G-F^* (APP KI) mice (n=7) and age-matched C57BL/6J control mice (n=8) as they explored familiar and novel contexts in the recording block. The percentage power values for oscillatory frequency bands were calculated and compared across genotype and recording sessions using a repeated measures Two-Way ANOVA. No significant main or interaction effects were detected at any frequency band. ***A*.** Delta (1-3 Hz): (Session x Genotype) F(3,39) = 2.735, *p*>0.05. (Session) F(2.053,26.69) = 2.093, *p*>0.05. (Genotype) F(1,13) = 0.841, *p*>0.05. ***B*.** Theta (4-12 Hz): (Session x Genotype) F(3,39) = 0.577, *p*>0.05. (Session) F(1.408,18.30) = 1.158, *p*>0.05. (Genotype) F(1,13) = 0.224, *p*>0.05. ***C*.** Beta (13-20 Hz): (Session x Genotype) F(3,39) = 0.470, *p*>0.05. (Session) F(2.604,33.85) = 1.621, *p*>0.05. (Genotype) F(1,13) = 0.293, *p*>0.05. ***D*.** Low Gamma (35-55 Hz): (Session x Genotype) F(3,39) = 2.091, *p*>0.05. (Session) F(2.329,30.28) = 1.300, *p*>0.05. (Genotype) F(1,13) = 0.387, *p*>0.05. ***E*.** High Gamma (65-120 Hz): (Session x Genotype) F(3,39) = 1.413, *p*>0.05. (Session) F(1.837,23.89) = 0.511, *p*>0.05. (Genotype) F(1,13) = 3.237, *p*>0.05. Top line graphs and error bars in each panel represent group means ± standard error mean (SEM). Bottom line graphs in each panel depict individual frequency band values for each mouse tracked across sessions. Control, solid connecting lines. APP KI, hashed connecting lines.

**Supplemental Figure 4.**
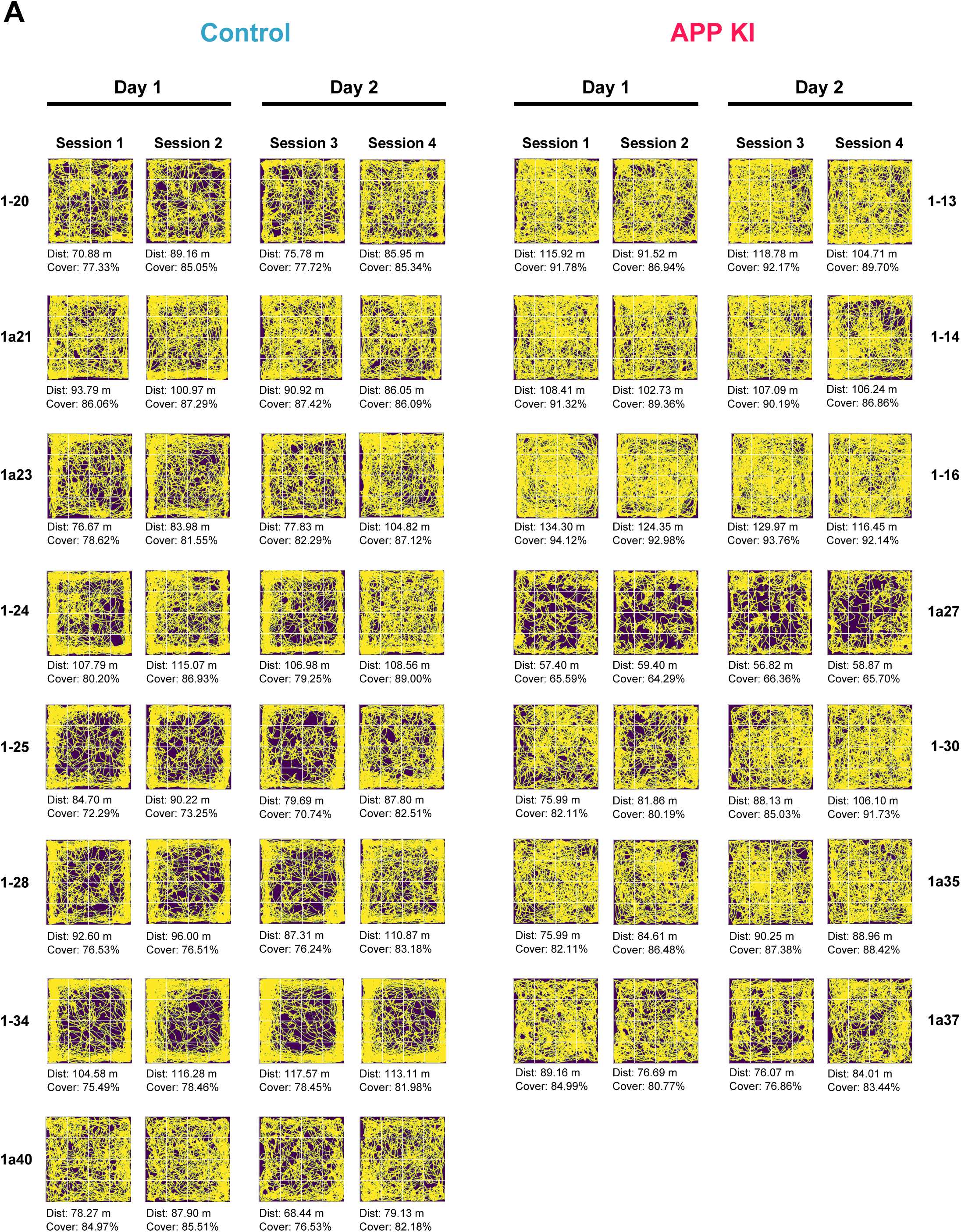
Recording block trajectory maps. ***A*.** Trajectory maps from every recording session in the recording blocks are shown for each mouse along with the total distance traveled (m) and % arena coverage. Mouse paths appear yellow on a dark background. A 4×4 grid (white lines) is overlaid onto the trajectory maps for Center Zone and Boundary Zone occupancy analysis appearing in Supplemental Figure 5. Arena dimensions: 50cm x 50cm.

**Supplemental Figure 5.**
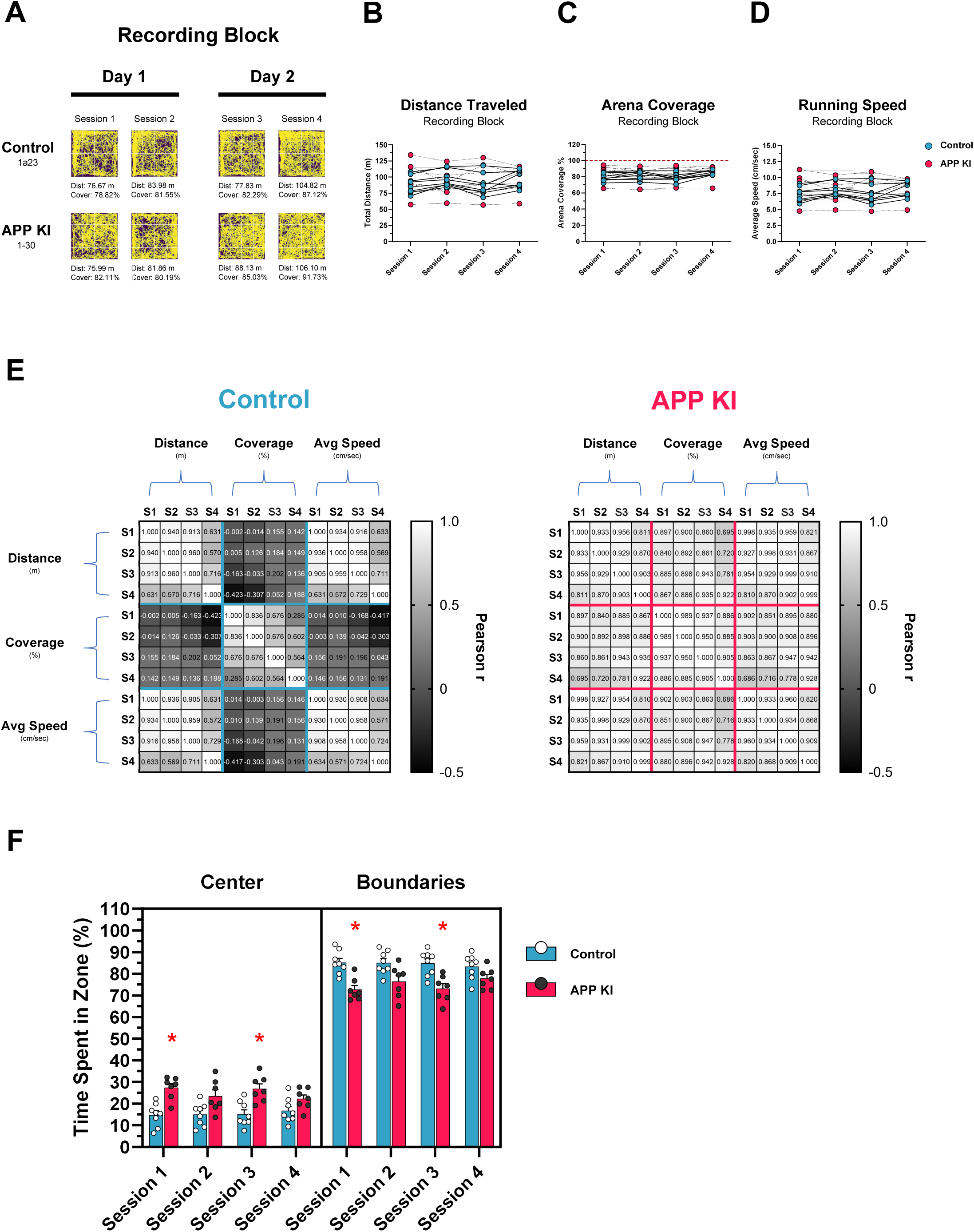
Aged *APP ^NL-G-F/NL-G-F^* mice exhibit increased exploration in the center zone of open field arenas in recording blocks. Behavioral performance for all 18-month mice through the recording paradigm was first assessed by calculating the total path length (m), percentage of arena coverage (%), and average speed (cm/sec) during exploration for each recording session. ***A*.** Representative trajectories for one mouse per group in the recording block are shown. Control, Mouse ID: 1a23; APP KI, Mouse ID: 1-30. Total distance traveled and % arena coverage values appear below each trajectory map. Trajectory paths appear yellow. ***B-D*.** Behavioral measures were tracked for each mouse in the recording block and analyzed using a repeated measures Two-Way ANOVA. No significant differences between genotype were detected on any measure. Repeated measures Two-Way ANOVA: Total distance traveled: (Session x Genotype) *F*(3,39) = 4.392, *p*<0.05. (Session) *F*(3,39) = 1.101, *p*>0.05. (Genotype) *F*(1,13) = 0.010, *p*>0.05. Arena Coverage: (Session x Genotype) *F*(3,39) = 4.815, *p*<0.01. (Session) *F*(3,39) = 4.812, *p<*0.05. (Genotype) *F*(1,13) = 1.046, *p*>0.05. Average speed: (Session x Genotype) *F*(3,39) = 4.532, *p*<0.01. (Session) *F*(3,39) = 1.069, *p>*0.05. (Genotype) *F*(1,13) = 0.025, *p*>0.05. ***E*.** To examine the relationship between total distance traveled, arena coverage, and average speed of our mice in the recording blocks, we generated correlation matrices comparing values for each genotype. All behavioral measures were found to be strongly correlated in APP KI mice across sessions, but not Controls. In Control mice, arena coverage was poorly correlated to distanced traveled and average speed. ***F*.** The 50cm x 50cm arenas were divided into 4×4 bins and assigned to one of two zones: Boundaries or Center. All bins along the arena borders were labeled Boundaries (12 bins) and remaining bins in the middle were labeled Center (4 bins). For each mouse and session, the total percentage time in each zone (%) was calculated and analyzed using a repeated measures Three-Way ANOVA. Each group spent significantly greater time exploring the boundaries of the arenas versus the center zone. However, APP KI mice explored the arena center more than Controls. This behavior was also reflected in the percentage time spent along the arena boundaries. Repeated measures Three-Way ANOVA: (Zone x Session x Genotype) *F*(3,39) = 2.920, *p*<0.05. (Zone x Session) *F*(3,39) = 1.100, *p*>0.05. (Zone x Genotype) *F*(1,13) = 15.100, *p*<0.01. (Session x Genotype) *F*(3,39) = 0.916, *p*>0.05. (Zone) *F*(1,13) = 583.000, *p*<0.001. (Session) *F*(3,39) = 0.167, *p*<0.05. (Genotype) *F*(1,13) = 5.230, *p*<0.05. Line graphs represent individual mouse performance values across recording block sessions. Bar graphs represent mean ± standard error mean (SEM) with individual mouse performance values overlaid on the graph. * *p*<0.05; ** *p*<0.01.

**Supplemental Figure 6.**
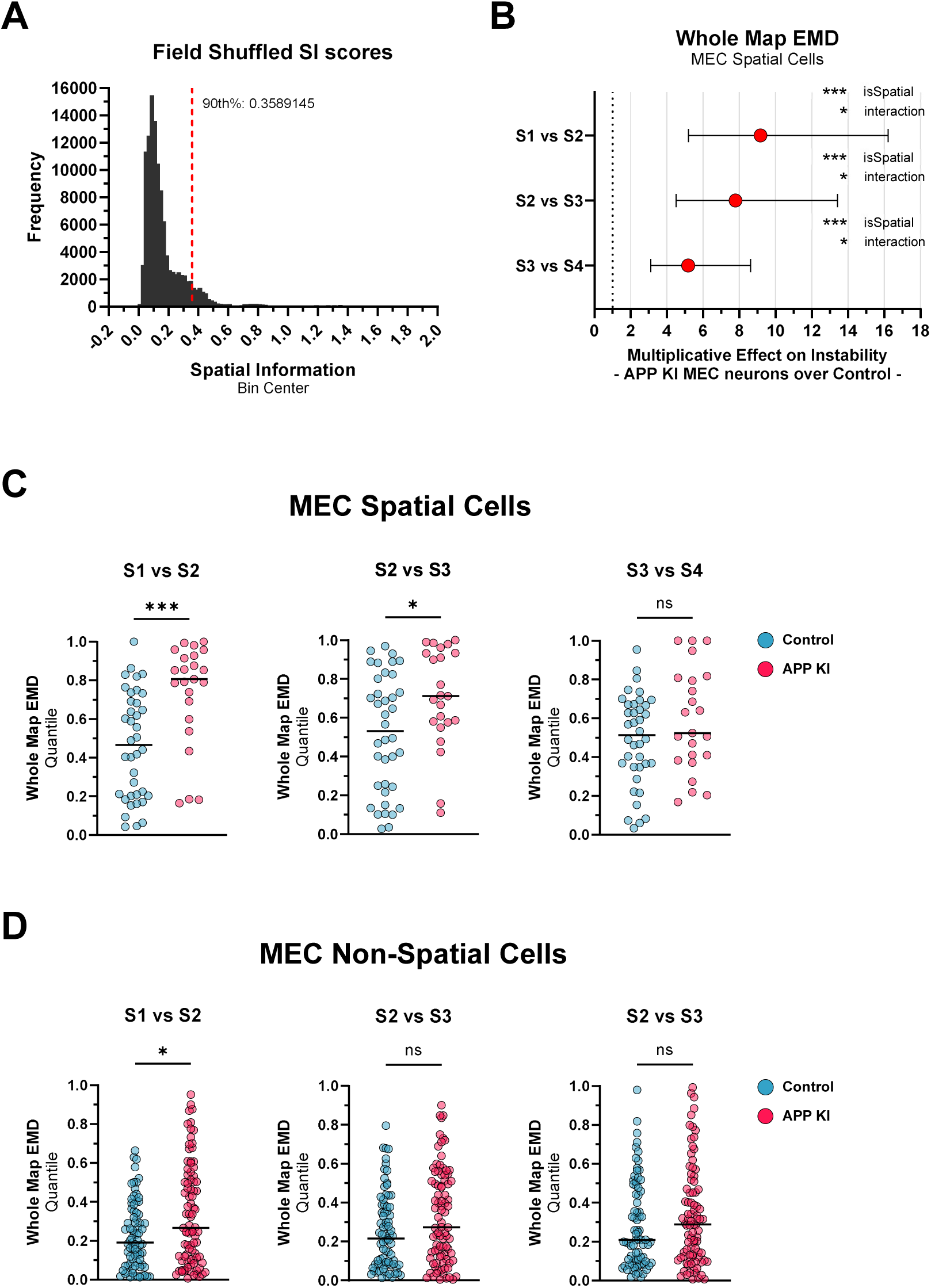
MEC spatial cells in aged *APP ^NL-G-F/NL-G-F^* mice identified using an alternative field shuffling procedure exhibit increased representational instability in familiar environments. In Figure 3, medial entorhinal cortex (MEC) spatial cells were identified from a distribution of shuffled spatial information (SI) scores generated by a circular shuffling procedure. This procedure misaligned spike times from *x*,*y* positions by circularly shifting spike timestamps by a constant between 20 seconds and T-20 seconds, where T is the duration of the recording session. As an alternative to this method of circular shuffling, we randomly shuffled the positions of segmented firing fields within their respective ratemaps using a procedure described here (Grieves *et al*., 2021). ***A*.** 1,000 field shuffles were performed for each Control MEC unit in Session 1 of our recording block and a distribution of field shuffled SI scores was created (Total, n=115,000 field shuffled SI scores). All units in Session 1 with true SI scores that exceeded the 90^th^ percentile of this distribution were classified as MEC spatial cells (90^th^ percentile: 0.3589 SI) (Control: n=38 spatial units, n= 77 non-spatial units; APP KI: n=23 spatial units, n=88 non-spatial units). Two APP KI units that were identified as spatial units after circular shuffling did not meet the 90^th^ percentile of field shuffled SI scores and were classified as non-spatial (APP KI: 1-14, T1C2; 1a35, T2C5). ***B*.** Whole map EMD analysis was run as described for Figure 3, with new spatial cell identification after field shuffling. A mixed-effects beta regression model found significant interaction effects of genotype and isSpatial variables for each session comparison (S1 vs S2, OR=9.17, *p*<0.05; S2 vs S3, OR=7.78, *p*<0.05; S3 vs S4, OR=5.18, *p*<0.05). The isSpatial variable had a significant main effect on all session comparisons (S1 vs S2, *p*<0.001; S2 vs S3, *p*<0.001; S3 vs S4, *p*<0.001), while genotype did not (S1 vs S2, *p*>0.05; S2 vs S3, *p*>0.05; S3 vs S4, *p*>0.05). ***C*.** Whole map EMD quantile values for MEC spatial cells identified by field shuffling were compared between genotype for each session comparison using two-tailed Mann-Whitney *U* tests. Quantile values were significantly larger in APP KI MEC spatial cells versus Control spatial cells in S1 vs S2 (*U*=195.5, *p*<0.001) and S2 vs S3 (*U*=272, *p*<0.05), but not in S3 vs S4 (*U*=347, *p*>0.05) comparisons. ***D*.** Quantile values of MEC non-spatial cells were also compared across genotype. Whole map EMD quantile values were significantly larger in APP KI MEC non-spatial cells versus Control non-spatial cells on S1 vs S2 (*U*=2,647, *p*<0.05) only. No genotype difference in quantile values were detected on S2 vs S3 (*U*=2,805, *p*=0.057) or S3 vs S4 (*U*=3,058, *p*>0.05) comparisons. For EMD analysis, ratemap bins were fixed across all animals and sessions (32X32 bins). EMD, Earth Mover’s Distance. OR, Odds Ratio. Purple datapoints in EMD OR plots indicate OR values with full range confidence intervals. Scatter plots represent median with full data range. * *p*<0.05; *** *p*<0.001. ns, not significant.

**Supplemental Figure 7.**
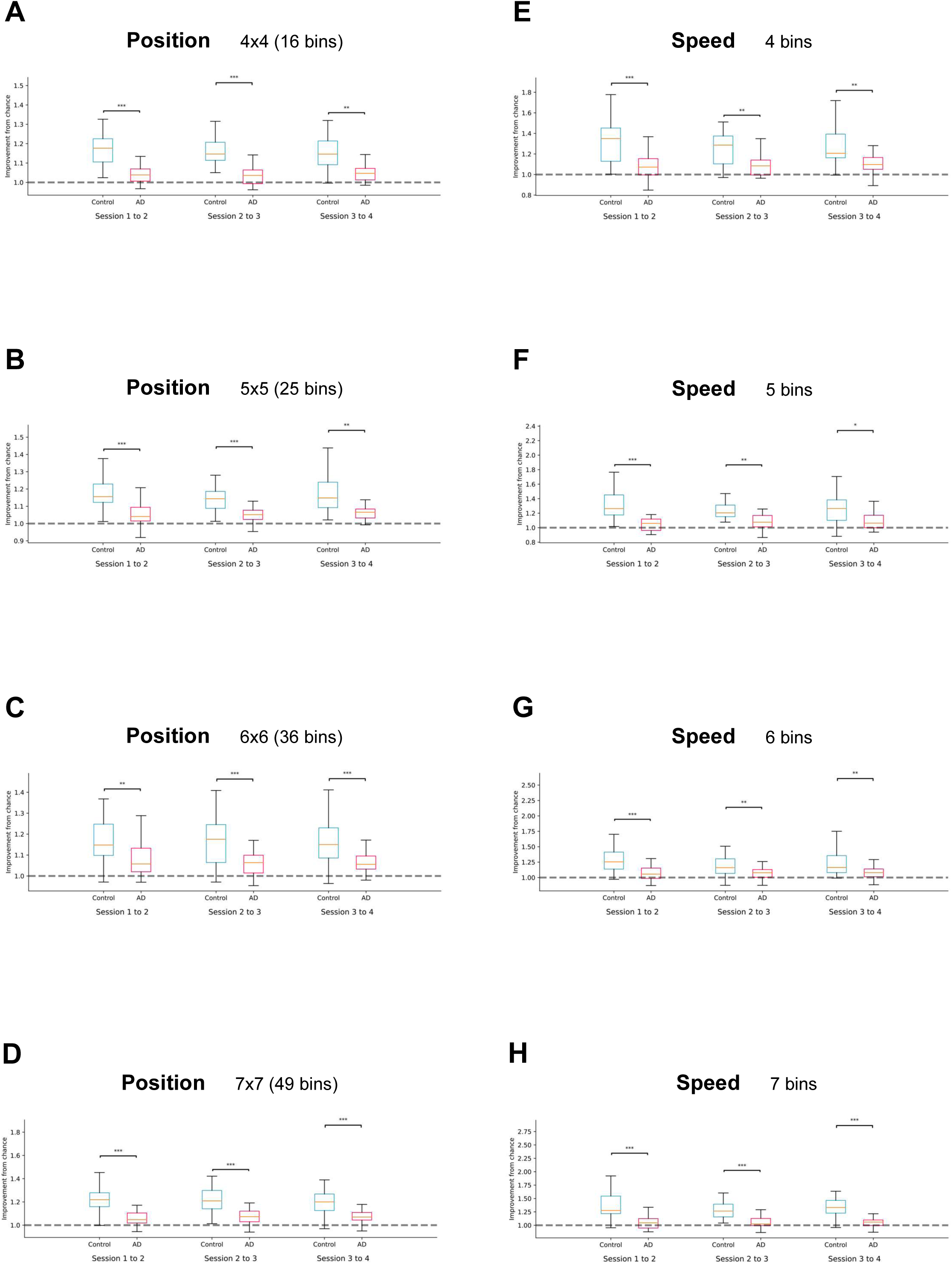
18-month *APP ^NL-G-F/NL-G-F^* mice exhibit impaired stability and generalization of position and speed coding in the medial entorhinal cortex. Linear support vector machines (SVMs) were used to decode discretized position and speed from a pooled set of neurons from aged mice in each group (APP KI vs Control). For cross-session analysis of data collected in our recording block, SVMs were trained on the full first session and tested in the second session for each session comparison (Session 1 vs Session 2, Session 2 vs Session 3, and Session 3 vs Session 4). For position, the arena space (50cm x 50cm) was divided into 16, 25, 36, and 49 spatial bins for individual runs of the analysis. For speed, the minimum-maximum speed distribution (3-100 cm/sec) was divided into 4, 5, 6 and 7 speed bins for individual runs of the analysis. ***A-D*.** Wilcoxon signed-rank tests were used to detect group differences in median decoding performance for position across sessions, expressed as an improvement from chance level. In all run conditions (spatial bins), MEC neurons from APP KI mice exhibited poor position decoding in each session comparisons versus Control. ***E-H*.** Group differences in median decoding performance for speed were detected in all run conditions (speed bins) and in all session comparisons using Wilcoxon signed-rank tests. MEC neurons from APP KI mice exhibited poor speed decoding for all session comparisons relative to Controls. Total session duration: 20 min. Box plot and whiskers represent the interquartile range (IQR) and median improvement from chance values per genotype for each test half-session comparison. Panels B and F appear in main Figure 4. * *p* < 0.05; ** *p* < 0.01; *** *p* < 0.001.

**Supplemental Figure 8.**
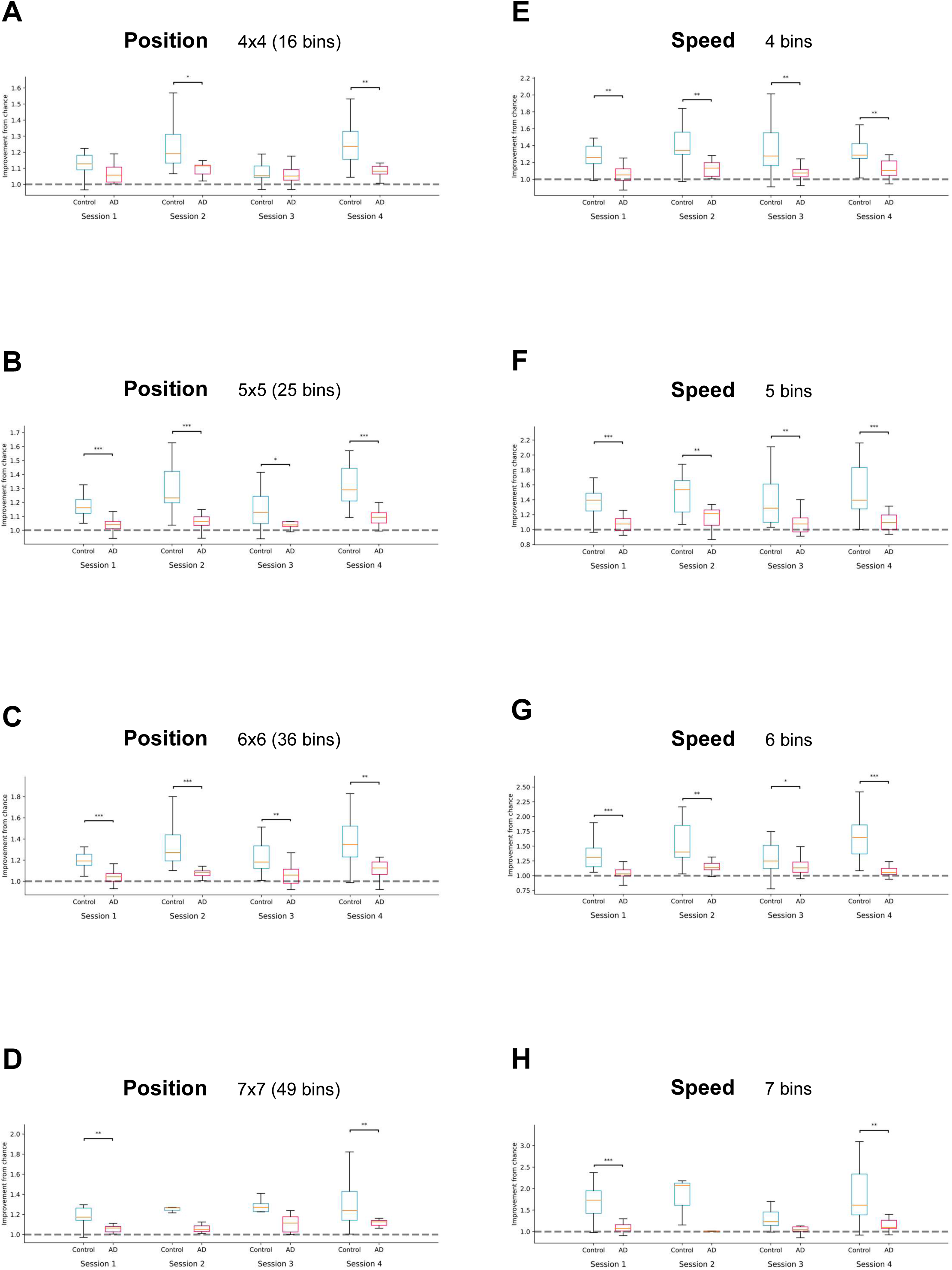
Medial entorhinal cortex neurons from aged *APP ^NL-G-F/NL-G-F^* mice exhibit impaired spatial coding within individual sessions of the recording block. Linear support vector machines (SVMs) were used to decode discretized position and speed from a pooled set of neurons for 18-month-old animals in each group (APP KI vs Control). For within-session analysis of data collected in our recording block, each recording session was split in half and the SVMs were trained on the first half and assessed on the second half. For position, the arena space (50cm x 50cm) was divided into 16, 25, 36, and 49 spatial bins for individual runs of the analysis. For speed, the minimum-maximum speed distribution (3-100 cm/sec) was divided into 4, 5, 6 and 7 speed bins for individual runs of the analysis. ***A-D*.** Wilcoxon signed-rank tests were used to detect group differences in median decoding performance for position, expressed as an improvement from chance level, in the tested half of each session. For runs of 25 and 36 spatial bins, MEC neurons from APP KI mice exhibited poor position decoding in each session of the recording block versus Control. ***E-H*.** Group differences in median decoding performance for speed were detected in the tested half of each recording session for runs of 4, 5, and 6 speed bins. For these conditions, MEC neurons from APP KI mice exhibited poor speed decoding performance within each session relative to Controls. Total session duration: 20 min. Half-session duration: 10 min. Box plot and whiskers represent the interquartile range (IQR) and median improvement from chance values per genotype for each test half-session comparison. * *p* < 0.05; ** *p* < 0.01; *** *p* < 0.001.

**Supplemental Figure 9.**
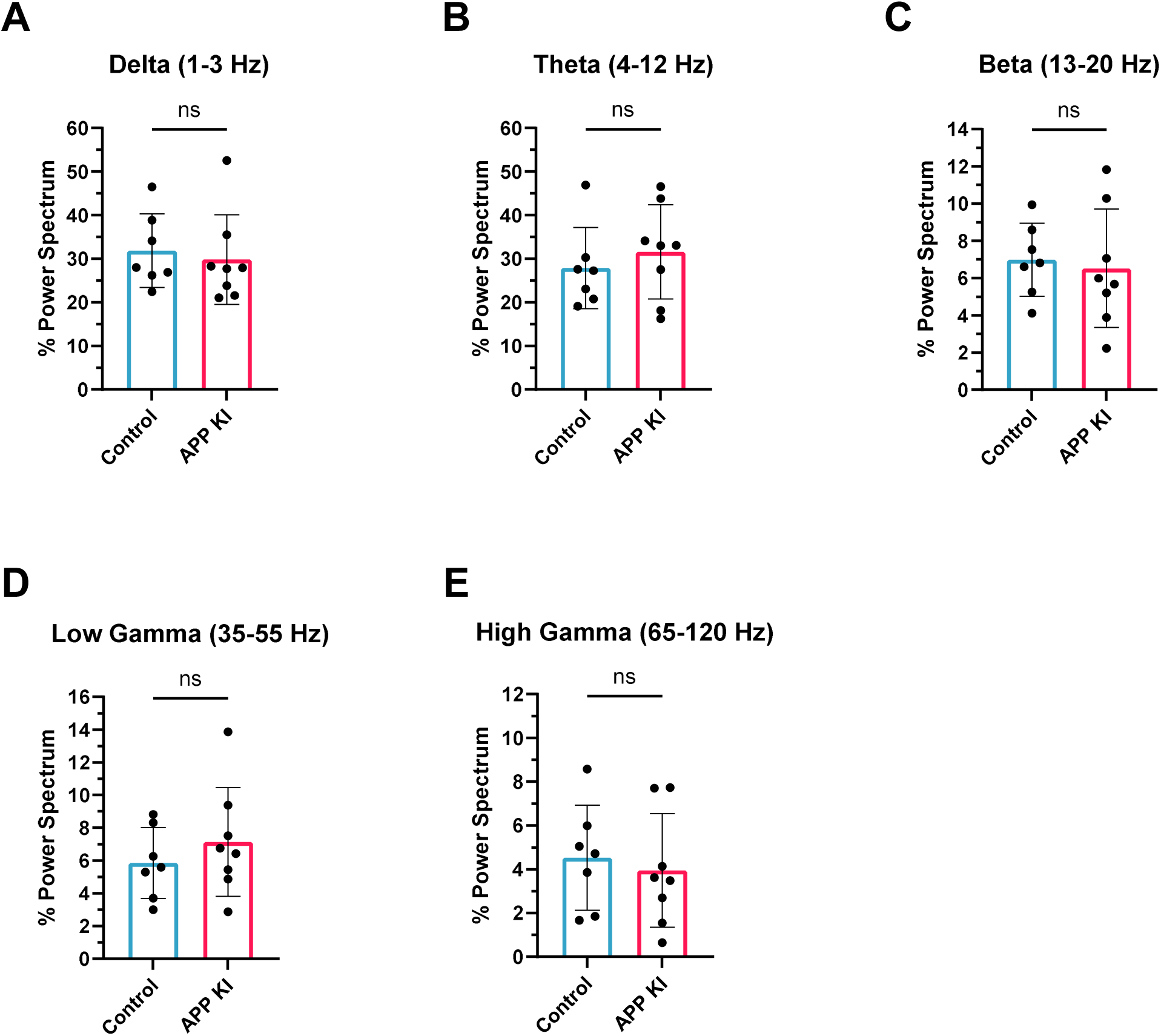
No effect of genotype on *in vivo* oscillatory activity in the medial entorhinal cortex of aged experimental mice. Local field potentials (LFP) were sampled in the medial entorhinal cortex (MEC) of 18-month *APP ^NL-G-F/NL-G-F^* (APP KI) mice and age-matched C57BL/6J control mice. The percentage power values for oscillatory frequency bands were first calculated for individual samples collected from multiple Session 1 recordings across different recording blocks per mouse (see Supplemental Data 1). An average % power spectrum value for each frequency band was then calculated per mouse. Finally, unpaired t-tests were used to compare the averaged % power spectrum values for each frequency band across genotype. No significant group differences were detected at any frequency band. ***A*.** Delta (1-3 Hz): *t*(13) = 0.4167, *P* = 0.6837. ***B*.** Theta (4-12 Hz): *t*(13) = 0.7066, *P* = 0.4923. ***C*.** Beta (13-20 Hz): *t*(13) = 0.3337, *P* = 0.7439. ***D*.** Low Gamma (35-55 Hz): *t*(13) = 0.8722, *P* = 0.3989. ***E*.** High Gamma (65-120 Hz): *t*(13) = 0.4494, *P* = 0.6606. Bar graphs represent mean ± standard deviation (SD). NS, not significant.

**Supplemental Figure 10.**
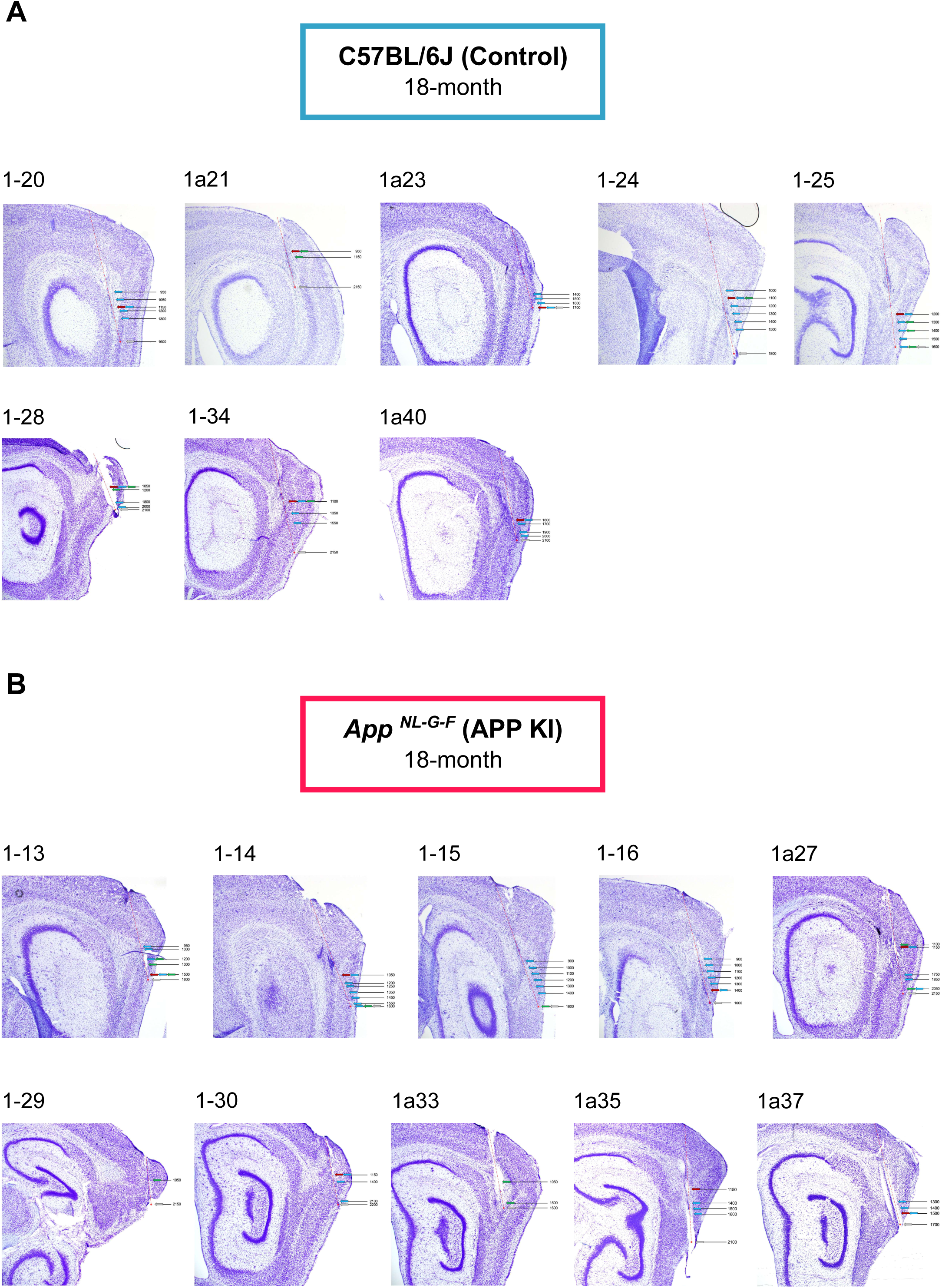
Tetrode trajectories and recording positions in aged *APP ^NL-G-F/NL-G-F^* mice and C57BL/6J controls. Histological verification of recording positions in 18-month C57BL/6J control mice (***A***) and age-matched APP KI mice (***B***) used in these studies. Brightfield images of cresyl violet stained sagittal sections appear for each mouse and show electrode trajectories (red dashed line). Red arrows indicate recording sites where units were collected for recording block analyses. Blue arrows indicate recording sites where units were sampled and pooled per mouse for hyperactivity analysis. Green arrows indicate recording sites where spatial cells were sampled. Grey arrows represent the last recording position for each mouse. Scale bars, 500µm.

**Supplemental Table 1.** Summary statistics for mixed-effects beta regression models following alternative field shuffling. The table shows the genotype (treatment), isSpatial, and interaction estimates and associated *p* values for each session comparison in Supplemental Figure 6. AIC and BIC values are also shown. Values correspond to Supplemental Figure 6B. AIC, Akaike Information Criteria. BIC, Bayesian Information Criterion. EMD, Earth Mover’s Distance.

| Dependent variable | Session comp. | AIC | BIC | Treatment estimate | Treatment *p* value | isSpatial estimate | isSpatial *p* value | Interaction estimate | Interaction *p* value |
| --- | --- | --- | --- | --- | --- | --- | --- | --- | --- |
| EMD quantile | S1 vs S2 | -109.1372 | -88.6140 | 0.3681 | 0.0881 | 1.0579 | 1.4981 x 10<sup>-7</sup> | 0.7896 | 0.0100 |
| EMD quantile | S2 vs S3 | -99.2234 | -78.7001 | 0.2229 | 0.2639 | 1.0551 | 1.7026 x 10<sup>-7</sup> | 0.7738 | 0.0118 |
| EMD quantile | S3 vs S4 | -65.5817 | -45.0585 | 0.1522 | 0.3769 | 0.7226 | 4.6164 x 10<sup>-4</sup> | 0.7698 | 0.0150 |

## Notes

### Competing Interest Statement

The authors have declared no competing interest.

